# Exploring Metabolic Anomalies in COVID-19 and Post-COVID-19: A Machine Learning Approach with Explainable Artificial Intelligence

**DOI:** 10.1101/2024.04.15.589583

**Authors:** Juan José Oropeza-Valdez, Cristian Padron-Manrique, Aarón Vázquez-Jiménez, Xavier Soberon, Osbaldo Resendis-Antonio

**Affiliations:** Human Systems Biology Laboratory. Instituto Nacional de Medicina Genómica (INMEGEN). Mexico City, Mexico; Centro de Ciencias de la Complejidad, Universidad Nacional Autónoma de México (UNAM). Mexico City, Mexico; Programa de Doctorado en Ciencias Biomédicas, Universidad Nacional Autónoma de México (UNAM). Mexico City, Mexico; Departamento de Ingeniería Celular y Biocatálisis, Instituto de Biotecnología, UNAM, México, Avenida Universidad 2001, Colonia Chamilpa, 62210, Cuernavaca, México; Coordinación de la Investigación Científica – Red de Apoyo a la Investigación, Universidad Nacional Autónoma de México (UNAM). Mexico City, Mexico

## Abstract

The COVID-19 pandemic, caused by SARS-CoV-2, has led to significant challenges worldwide, including diverse clinical outcomes and prolonged post-recovery symptoms known as Long COVID or Post-COVID-19 syndrome. Emerging evidence suggests a crucial role of metabolic reprogramming in the infection’s long-term consequences. This study employs a novel approach utilizing machine learning (ML) and explainable artificial intelligence (XAI) to analyze metabolic alterations in COVID-19 and Post-COVID-19 patients. By integrating ML with SHAP (SHapley Additive exPlanations) values, we aimed to uncover metabolomic signatures and identify potential biomarkers for these conditions. Our analysis included a cohort of 142 COVID-19, 48 Post-COVID-19 samples and 38 CONTROL patients, with 111 identified metabolites. Traditional analysis methods like PCA and PLS-DA were compared with advanced ML techniques to discern metabolic changes. Notably, XGBoost models, enhanced by SHAP for explainability, outperformed traditional methods, demonstrating superior predictive performance and providing different insights into the metabolic basis of the disease’s progression and its aftermath, the analysis revealed several metabolomic subgroups within the COVID-19 and Post-COVID-19 conditions, suggesting heterogeneous metabolic responses to the infection and its long-term impacts. This study highlights the potential of integrating ML and XAI in metabolomics research.

## Introduction

The COVID-19 pandemic, caused by the coronavirus SARS-CoV-2, has presented a formidable challenge to global health systems. As of March 2024, the number of confirmed COVID-19 cases has surpassed 770 million ^1^. The wide spectrum of symptoms, varying from mild to severe respiratory distress and multi-organ dysfunction ^2^, underscores the need for a comprehensive systemic understanding of the disease’s pathophysiology and the factors contributing to its diverse clinical outcomes ^3,4^. In addition to the immediate health impacts, the COVID-19 pandemic has highlighted the long-lasting effects and challenges in the post-recovery phase. Many individuals who have recovered from COVID-19 have reported a wide range of persistent symptoms and health issues ^5,6^. Common symptoms following recovery include persistent fatigue, shortness of breath, cough, joint and chest pain, brain fog, depression, and anxiety ^7,8^. Moreover, the full extent of these symptoms and their long-term consequences remain uncharacterized. The post-recovery symptoms, often referred to as “Post-COVID-19” or “Long COVID-19” syndrome, can persist for weeks or up to 2 years after the initial infection ^9^. Although certain mechanisms, viral persistence ^10^, immune dysregulation ^7^, and organ damage ^11^, have been identified as potentially involved in Post-COVID-19 symptoms, their exact understanding remains incomplete. One suitable factor accompanying the post-symptoms is metabolic reprogramming at the systemic level. Emerging evidence suggests these long-term consequences may be linked to systemic metabolic reprogramming during infection, affecting pathways related to amino acids, glucose, cholesterol, fatty acids, among others^12^. This metabolic disruption alters energy production and immune regulation, pointing to a need for further research to understand these changes and develop specific therapeutic interventions.

Metabolomics offers a comprehensive and unbiased view of the biochemical alterations occurring during viral infections portraying the complex interactions between the viral pathogen and the host response ^13,14^. Notably, this approach has been proven successful in uncovering distinct metabolic signatures associated with various infectious diseases ^15^, including COVID-19^16^ and Post-COVID-19^17,18^. However, metabolomics profiles have high dimensionality nature^19^ and can exhibit nonlinear interactions. Traditionally, linear dimensionality reduction methods are used to identify low-dimensional embedding spaces in metabolomic data. Among these methods, PCA (Principal Component Analysis) and its supervised counterpart the PLS-DA (Partial Least Square Discriminant Analysis) ^20^ are the most frequent. Despite their importance, these methods exhibit significant limitations when it comes to uncovering and analyzing nonlinear interactions, which are often crucial in differentiating intricate groups, such as control versus disease phenotype^21^. Alternatively, differential expression analysis applied in metabolic concentrations is a well-established technique to identify metabolites with significant statistical differences expressed between clinical groups. This latter strategy solely captures the over-representation of features within a class identified by the magnitudes of these changes using p-values. As a result, it falls short in detecting complex interactions. To overcome this limitation, some non-supervised and supervised machine learning algorithms have been suggested to take into account the linear and non-linear interactions emerging from metabolome data. For instance, Uniform Manifold Approximation and Projection (UMAP), an unsupervised reduction method in multidimensional data, captures the complex topology of high-dimensional spaces and effectively reduces it to a lower-dimensional representation. This approach provides superior projections and enhanced cluster separation in handling intricate data structures compared to other dimensional reduction methods such as PCA, t-SNE, and even autoencoders^22^. However, features with low variable magnitude typically have a reduced impact on these low-dimensional projections due to their dependence on distance metrics, even though they can be informative for phenotype classification. Furthermore, supervised machine learning algorithms, like eXtreme Gradient Boosting (XGBoost) ^23^, have emerged as a solution to identify those variables that play an important role in classifying groups of multidimensional data. This classification algorithm is insensitive to feature magnitude variations, capable of discerning subtle and or complex patterns, transcending the limitations of traditional methods and over-representation biases ^23,24^. The insensitivity of XGBoost to feature magnitude means that it does not require extensive data preprocessing to normalize or to scale the features, making it more robust and easier to apply due to its tree-based method. ^23,24^. Notably, by combining this approach with the SHAP (SHapley Additive exPlanations) method, XGBoost goes beyond detecting the high and low magnitudes of metabolites to classify a phenotype, particularly, the SHAP (SHapley Additive exPlanations) values provide a metric to determine the unique contributions of individual features. SHAP values can either be used to rank the importance of each feature in making predictions (global explainability) or to elucidate how the individual predictions are derived (local explainability) ^25^. Interestingly, local explainability can be employed for supervised clustering to create explainable embeddings^25^. So, for a metabolome dataset, each sample’s complex, multi-dimensional metabolic profile can be represented in a reduced dimensional space while preserving the explainability of individual features for the prediction^26^. As a remarkable property, these explainable embeddings space is unbiased by magnitude or scale of the variables when we used XGBoost. In this type of explainable embedding, similarities between samples are determined by the importance weight for classification rather than the original values^23,25^. While explainable embeddings have been employed in metabolomics studies^27^, they have never been used before in COVID-19 or Post-COVID-19 studies. Therefore, there is a need to use these new approaches to identify novel groups related to Post-COVID-19, particularly in areas where traditional unsupervised methods reach their limits.

In the context of understanding metabolic anomalies between COVID-19 and Post-COVID-19 phenotypes, the objective of the study is to contrast the biomarkers obtained from previous statistical techniques already published ^16,28^ with advanced machine learning algorithms combined with analyses of global and local explainability. Altogether, our study allowed us for a detailed and multifaceted exploration of identified metabolites. Our analysis not only suggests potential biomarkers through differential expression analysis but also points to the complexity of metabolic alterations through subgroup analysis and local explainability combining machine learning and XAI.

## Results

### Overview of the analysis and cohort study

With the aim of extending the list of metabolites, and far away from those identified by linear methods, that serve as biomarkers to differentiate normal, COVID-19, Post-COVID samples, we implemented some machine learning algorithms. Figure 1 illustrates a comprehensive workflow of this study’s analytical process, breaking it down into traditional analysis and machine learning approaches. The metabolomics data were obtained from previous reports freely available in these references ^16,28^. In summary, data consists of 111 identified metabolites across three classes: 142 COVID samples, 48 post-COVID samples, and 38 control samples from published datasets (See methods: Data). Our analytical workflow is divided into three main branches. In the first one, we combine classical linear and nonlinear dimensionality reduction methods to explore potential features differentiating each clinical group. Thus, dimensional reduction techniques such as PCA (unsupervised), PLS-DA (supervised), and UMAP (unsupervised) are applied to the data in this section. Additionally, we conducted differential expression analysis using Earth Mover’s Distance (EMD), as a complementary strategy to identify over-represented markers. The second branch is devoted to implementing some supervised machine learning algorithms to classify the clinical data. To construct a computational model with the best predictable capacity, we assessed four methods of classification (Logistic Regression, Support Vector Machine, Random Forest, and XGBoost) and selected the one that had the best performance among them. Once selected the model with the best performance, we carefully and extensively survey the importance of the global explainability of each feature through the application of SHAP. The calculation of SHAP values offers a means to interpret the model by assigning a mean weight of feature importance that is not biased by the scale of the data.

**Figure 1.**
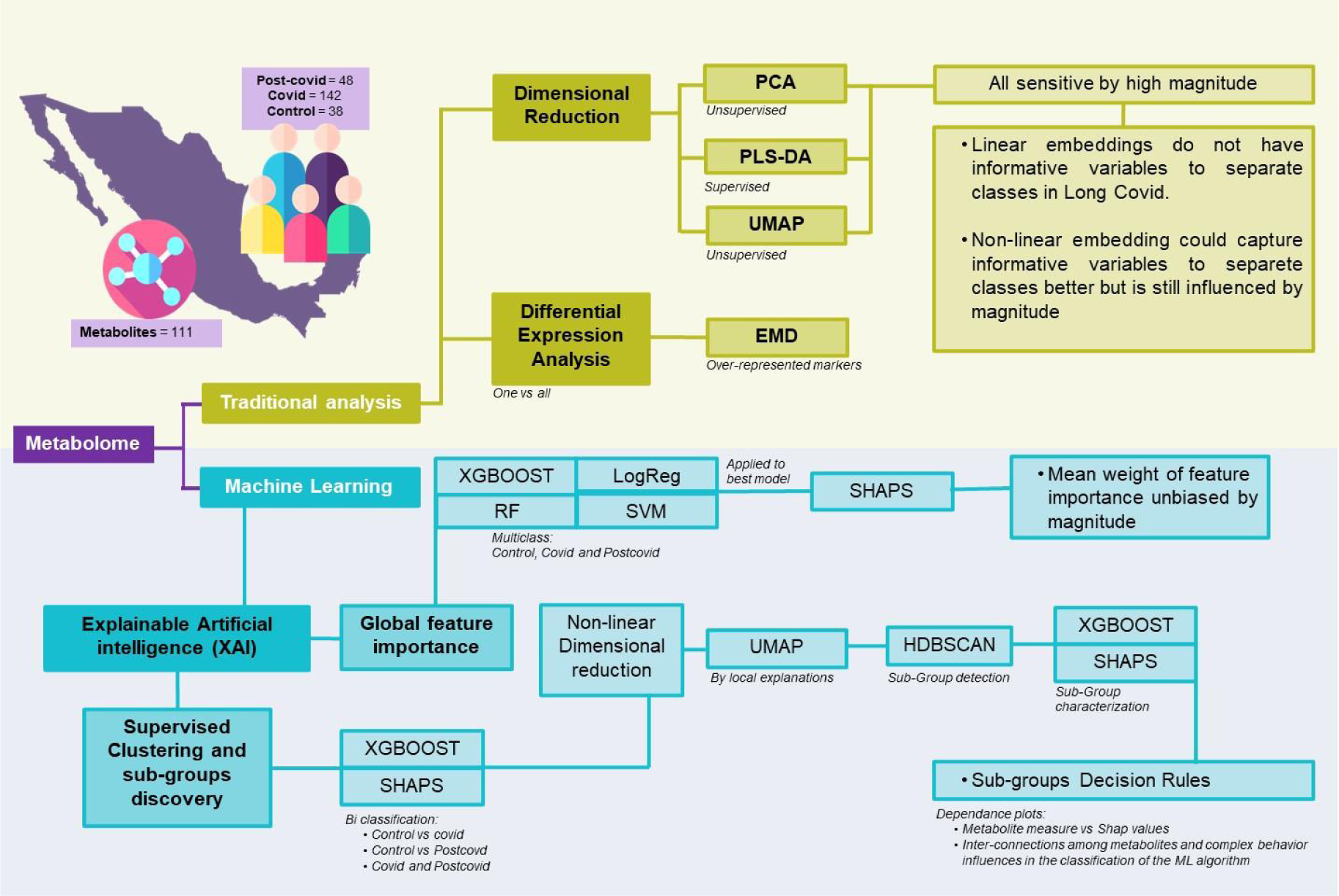
Schematic representation of various analytical approaches applied to the integrated metabolome data. The acronyms and their respective meanings are: **PCA**: Principal Component Analysis - An unsupervised method that transforms the original variables into a new set of variables called principal components. **PLS-DA**: Partial Least Squares Discriminant Analysis - A supervised technique that seeks to find a linear combination of features that best separates two or more classes in a dataset. **UMAP**: Uniform Manifold Approximation and Projection - A non-linear dimensionality reduction technique that works well for clustering and visual representation of high-dimensional datasets. **EMD**: Earth Mover’s Distance - A measure of the distance between two probability distributions, conceptualized as the minimum “work” needed to transform one distribution into another. It is also employed as a measure for differential expression by clusters. **XAI**: Explainable Artificial Intelligence - A branch of AI that aims to make the decision-making process of machine learning models transparent and understandable. **XGBOOST**: Extreme Gradient Boosting - A highly efficient and scalable implementation of gradient boosting that works for both regression and classification problems. **RF:** Random Forest - An ensemble method that builds multiple decision trees for robust classification and regression outputs. **SVM:** Support Vector Machine - A powerful classifier that finds the optimal hyperplane for categorizing data into two distinct classes. **LogReg:** Logistic Regression - A statistical model that estimates probabilities of binary outcomes based on input features, adaptable to multiclass problems. **SHAPS**: SHapley Additive exPlanations - A method to explain individual predictions of any machine learning model by computing the contribution of each feature to every prediction. **HDBSCAN**: Hierarchical Density-Based Spatial Clustering of Applications with Noise - An advanced clustering algorithm that identifies clusters of varying shapes and sizes from a dataset.

The third approach focuses on nonlinear dimensionality reduction and clustering analysis to explore the characterization and local explainability of the data. To achieve this goal, we proceeded as follows. Starting from the model with the best performance, we trained it through binary classification between pair conditions and accomplished a post-hoc analysis using SHAP values. Afterwards, we utilized nonlinear dimensionality reduction via UMAP to elucidate local explanations, providing insights into the formation of subgroups within the high-dimensional SHAP values data. To identify samples containing a set of metabolites with similar weights of classification for each subgroup, we applied the Hierarchical Density-Based Spatial Clustering of Applications with Noise algorithm (HDBSCAN). Having identified each subgroup, the final step involved formulating decision rules for each of them. To this end, we conducted a multi-class classification of these clusters with XGBoost and obtained their SHAP values. To understand the specific decision rules for each subgroup we used the dependency plot (SHAP value vs original magnitude, for example see Supplementary Figure 1). In the next sections, we present the results obtained for each analysis.

### Limited Discrimination by Traditional Methods in Metabolic Profiling

Utilizing PCA, the inherent variance within the dataset was initially assessed. As displayed in Figure 2A, the CONTROL, COVID-19, and POST-COVID-19 samples exhibited overlapping regions, emphasizing the complexity of the metabolic patterns using this method alone. Despite PC1 accounting for 16.3% of the variance and PC2 capturing an additional 7.4%, these components did not offer a comprehensive separation of the groups. Similarly, the PLS-DA attempted to maximize the discrimination between the predetermined groups (Figure 2B). While it highlighted some tendencies, the results still showed overlaps, indicating that linear methods, such as PCA and PLS-DA, might not be sufficient to capture the intricate variations present in the metabolites. Different normalization/transformation strategies showed similar trends (Supplementary Figure 2)

**Figure 2.**
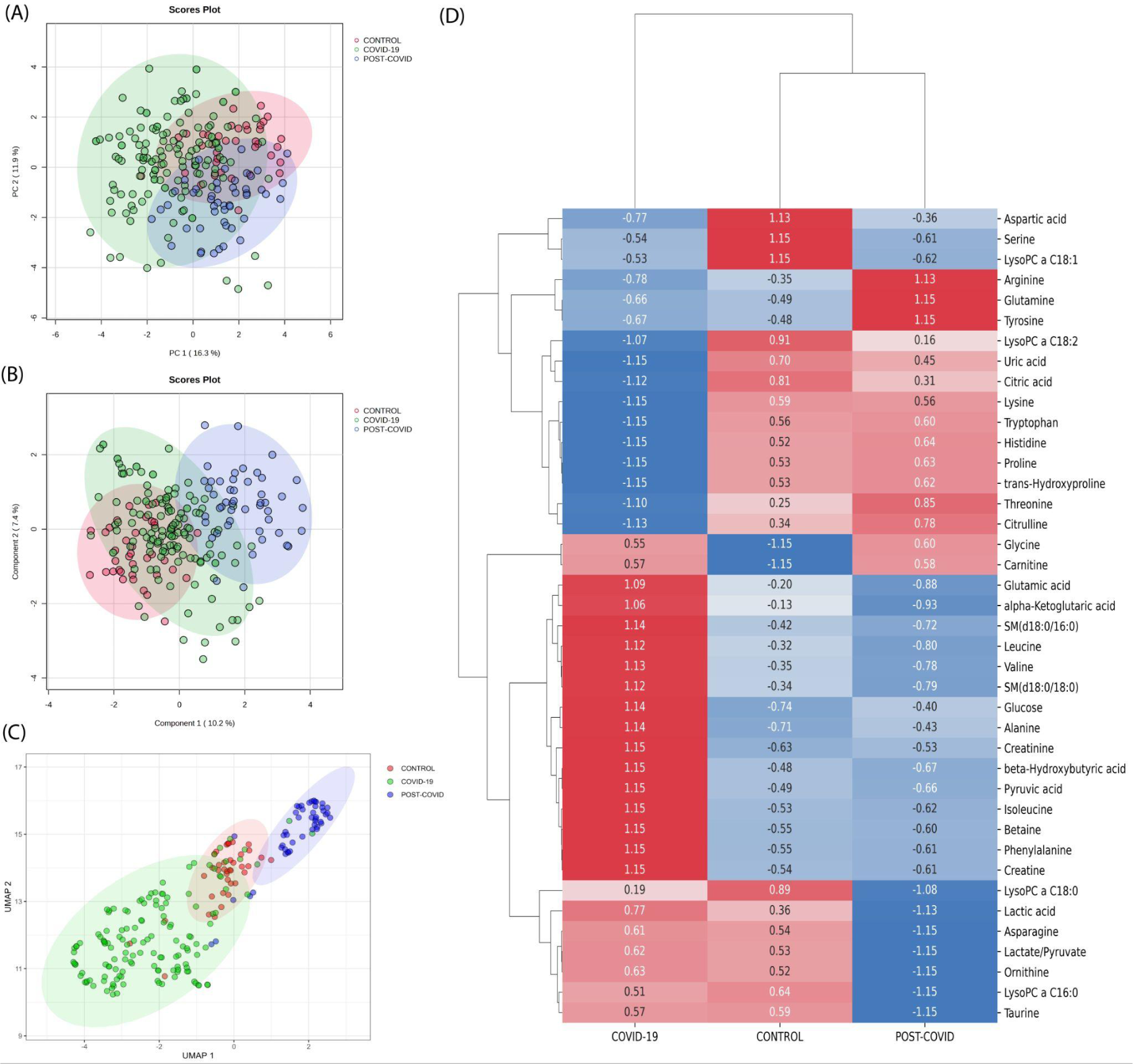
Traditional metabolic Analysis across CONTROL, COVID-19, and Post-COVID-19 States. **(A)** Principal Component Analysis (PCA) scores plot representing the metabolic profiles across samples, with the percentage of variance denoted by PC1 and PC2. **(B)** Partial Least Squares-Discriminant Analysis (PLS-DA) scores plot suggesting the metabolic tendencies of the CONTROL, COVID-19, and Post-COVID-19 groups, with variance represented by Component 1 and Component 2. For PCA and PLS-DA data median normalization, log transformation, and Pareto scaling were used. **(C)** Uniform Manifold Approximation and Projection (UMAP) visualization of metabolomic data across different conditions. Each point represents an individual sample. The color coding corresponds to the three conditions: CONTROL (green), COVID-19 (red), and POST-COVID-19 (blue). The distinct clustering of samples indicates metabolic differences among the groups **(D)** Heatmap derived from Earth Movers Distance (EMD) analysis illustrating the differential expression of metabolites across the CONTROL, COVID-19, and POST-COVID-19 groups. Displayed values represent the EMD, with positive numbers indicating a higher distribution cost relative to other groups, and negative numbers indicating a lower cost. Colors range from blue (lower EMD cost) to red (higher EMD cost), reflecting the magnitude and direction of metabolite level changes among the 40 most expressed metabolites.

On the other hand, we discerned distinct clustering patterns among the three groups (CONTROL, COVID-19, and POST-COVID-19) when one applied UMAP dimensional reduction (Figure 2C). With UMAP, data points representing the COVID-19 group predominantly occupied the lower left quadrant, exhibiting a more dispersed and non-linear distribution. In contrast, the CONTROL group’s data points seemed to concentrate around the center, exhibiting a tighter clustering pattern with sporadic overlap with the group, the POST-COVID-19 group manifested an elongated cluster formation extending towards the upper right quadrant. Notably, while there was some overlap between the COVID-19 and POST-COVID-19 groups, the latter’s data points were distinctly separate from the CONTROL group. This result suggests that non-linear projection could contribute to a better separation of the data.

Dimensionality reduction methods, including UMAP, are sensitive to high-abundance metabolites, which can overshadow less abundant ones, affecting low-dimensional embeddings. Therefore, class separation might be driven by a few dominant metabolites. For this reason, we employed differential expression analysis to identify high-abundance metabolites in class distinctions, offering a complementary analysis to dimensionality reduction methods.

### Differential Metabolite Expression using Earth Mover’s Distance (EMD)

Complementing our dimensionality reduction analyses, EMD (one vs all strategy) was utilized to capture the spectrum of metabolic variations, providing a measure of the distributional shifts between metabolites per conditions. EMD revealed distinct patterns of metabolite variations across the three conditions: CONTROL, COVID-19, and POST-COVID-19 (Figure 2D). In the CONTROL group, several metabolites, including aspartic acid, serine, and LysoPC a C18:1, were found to be more prevalent, as indicated by the positive EMD values. The COVID-19 group showed that Arginine and glutamine levels exhibited significantly lower levels of arginine and glutamine, which may reflect metabolic disturbances due to the viral infection. Citrulline and threonine also showed reduced levels in this group. POST-COVID-19 phase, the metabolite profile did not fully revert to that of the CONTROL group. Some metabolites, like proline and trans-hydroxyproline, approached the baseline levels observed in the CONTROL group, while others, such as glycine and carnitine, remained altered. Several metabolites clustered together in terms of their expression patterns, for instance, lactic acid, leucine, alpha-ketoglutaric acid, and glutamic acid showed a synchronous increase in the COVID-19 group and a subsequent decline in the POST-COVID-19 phase, potentially pointing towards a coordinated metabolic response or shared biochemical pathway. EMD captured metabolic differences that the linear analysis like PCA and PLS-DA, did not detect, it revealed distributional differences between conditions that informed on metabolites overlooked by linear analyses as shown in the limited intersections in Supplementary Figure 3.

Although the EMD matches dissimilarity between the metabolome distributions between groups ignoring if there is a linear or nonlinear dependency, it relies on the magnitude and dispersion on the metabolome distribution. This approach highlights the hidden information beneath the linear dependence space in which the PCA and PLS-DA stay. Moreover, to address class disparities that are not discernible through conventional methodologies sensitive to magnitude, it is imperative to integrate additional analytical strategies that are not magnitude-sensitive, such as ML approaches.

#### Evaluation of Multiclass Machine Learning Models and XAI

While our preceding analyses were insightful, they have the predisposition to emphasize features with higher/lower magnitudes, this can inadvertently overshadow subtler but crucial differences in the metabolites ^29^. Recognizing this limitation, we transitioned to machine learning (ML) models, aiming to harness their ability to predict and classify without unduly favoring dominant features. To this end, we employed 4 different machine learning algorithms, XGBoost, Random Forest (RF), Support Vector Machine (SVM), and Logistic Regression (LogReg) to identify metabolites whose concentrations can distinguish the physiological stages of the individuals. As shown in Figure 3, the XGBoost model had the highest predictive performance with a micro and macro Area Under the Curve (AUC) of 0.99 over other ML models showing excellent discrimination (Table 1).

**Figure 3:**
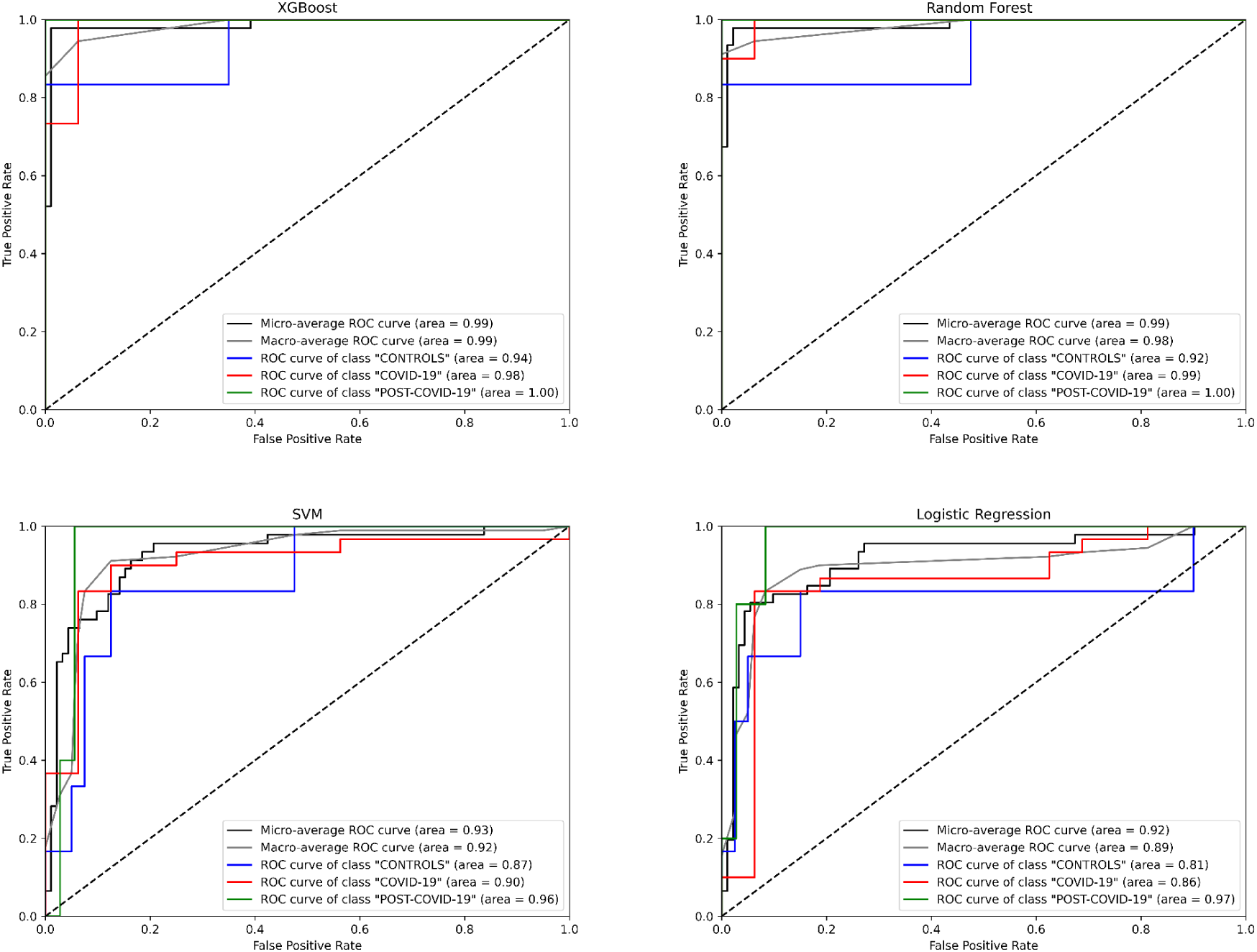
Receiver Operating Characteristic (ROC) curves for different machine learning models predicting three classes: “CONTROLs”, “COVID-19”, and “POST-COVID-19”. Four algorithms were evaluated: XGBoost, Random Forest, SVM and Logistic Regression. The curves depict the true positive rate (Sensitivity) against the false positive rate (1-Specificity) for each class. The diagonal dashed line represents the line of no-discrimination. AUC (Area Under the Curve) values are provided for micro and macro average ROC curves, as well as individual class ROC curves, Blue: CONTROLs, Red: COVID-19, Green: POST-COVID-19.

**Table 1.** Performance metrics of machine learning multiclass models. The mean and standard deviation of a variety of metrics of performance of each model are evaluated through cross-validation.

| ML Method | Train set + Cross Validation |  |  |  | Test Set |  |  |  |  |  |
| --- | --- | --- | --- | --- | --- | --- | --- | --- | --- | --- |
|  | Accuracy | Precision | Recall | F1 Score | Accuracy | Precision | Recall | F1 Score | Micro AUC | Macro AUC |
| XGBoost | 0.907 ± 0.059 | 0.920 ± 0.070 | 0.879 ± 0.073 | 0.885 ± 0.067 | 0.978 | 0.989 | 0.944 | 0.964 | 0.99 | 0.99 |
| **RF** | 0.891 ± 0.053 | 0.905 ± 0.077 | 0.848 ± 0.073 | 0.854 ± 0.067 | 0.957 | 0.979 | 0.911 | 0.941 | 0.99 | 0.98 |
| **SVM** | 0.819 ± 0.039 | 0.793 ± 0.042 | 0.785 ± 0.056 | 0.782 ± 0.045 | 0.804 | 0.716 | 0.7 | 0.707 | 0.93 | 0.92 |
| **LogReg** | 0.852 ± 0.071 | 0.867 ± 0.065 | 0.806 ± 0.106 | 0.808 ± 0.097 | 0.804 | 0.731 | 0.767 | 0.744 | 0.92 | 0.89 |

To describe our best model’s explainability, we explored the XGBoost model’s decision-making process by obtaining its SHAP values. Figure 4 provides a comprehensive SHAP analysis. Figure 4A underscores the overall influence of each metabolite in the model when classifying each physiological group. As the same figure shows, the most important variable to classify the groups are ratios Kynurenine/Tryptophan and Lactacte/Pyruvate, PC aa 36:6, Taurine, Glutamine, Phenylalanine, LysoPC a C26:0, Spermidine, Tryptophan, Glucose, LysoPC a C16:0 and Sarcosine emerging as top salient features. In Figure 4B, individual sample-level SHAP values are portrayed across three categories: COVID-19, CONTROLs, and Post-COVID-19. In each category, the XGBoost showed different important metabolites with its SHAP explanations. In CONTROLS, the top metabolites based on the SHAPs are ratio Kynurenine/Tryptophan, ratio Lactate/Pyruvate, LysoPC a C18:2, Glucose, Decadienylcarnitine, and Kynurenine. For COVID-19, the most influential metabolites to distinguish this class from the other are: PC aa C36:6, Spermidine, Tryptophan, Phenylalanine and ratios Kynurenine/Tryptophan, Lactate/Pyruvate. Lastly, for patients with POST-COVID-19 symptoms, the key metabolites differentiating from the other stages are Taurine, Glutamine, LysoPC a C16:0, Lactate/Pyruvate, and Sarcosine.

**Figure 4:**
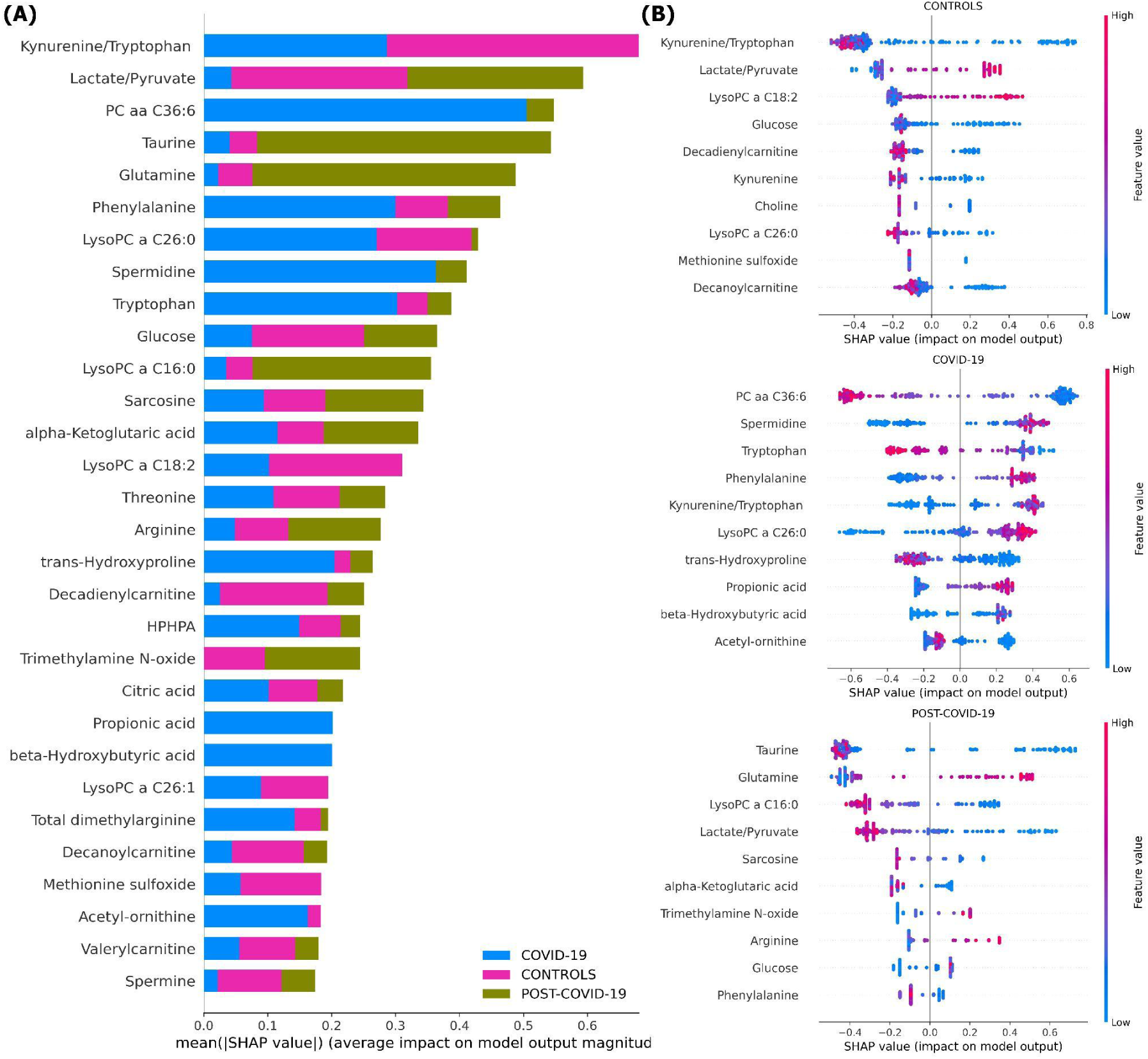
SHAP value analysis of an XGBoost model trained on metabolite data for multiclass classification. **A)** Average magnitude of SHAP values for each metabolite, indicating their overall importance in the model (Global explainability). **B)** Dot plots representing individual SHAP values (Local explainability) for each comparison following the Local explainability uses a ‘one vs all’ approach. For instance, at the top we show the Global explainability of the CONTROL vs the rest of the samples (COVID-19 and POST-COVID-19). Colors denote the relative concentration of metabolites, with blue indicating low and pink indicating high concentrations. For example, positive SHAP values for a sample (indicated by a dot) reflect the importance of a feature in classifying a specific category, such as distinguishing ‘CONTROL’ from other sample groups, including COVID-19 and POST-COVID-19.”

### Metabolomic Profiling and Binary Model Interpretation Using SHAP Values

To better understand how the best machine learning model algorithm (XGBoost) classifies each physiological group and find the most important metabolites that explain the classification, we proceed to built XGBoost models for all binary classification between pairs of conditions (CONTROL vs COVID-19, CONTROL vs POST-COVID-19 and COVID-19 vs POST-COVID-19). This strategy provided insights into the mean weight of feature importance (global explanations), asserting its robustness against data scale biases and the adoption of explainable artificial intelligence (XAI) techniques facilitated a more transparent interpretation of our machine learning models.

Performance metrics of the XGBoost models for each pairwise comparison showed an average of optimal classification similar to the multiclass model (Supplementary table 1). In Figure 5, we present the visualization of the SHAP values derived from binary XGBoost models for the three comparisons. Panel A depicts the SHAP values when comparing CONTROL to COVID-19 samples. Here, our model suggests that Phenylalanine, Kynurenine/Tryptophan, and Decadienylcarnitine showcase notable distinctions between the two groups. Similarly, Panel B depicts the SHAP values for CONTROL and POST-COVID-19 samples. In this case, LysoPC a C16:0, Glucose, Taurine, and the ratio Lactate/Pyruvate emerge as significant metabolites distinguishing these two groups. Lastly, Panel C offers a comparison between COVID-19 and POST-COVID-19 samples revealing metabolites like Glutamine/Glutamate, Taurine, Lactic acid, and alpha-ketoglutaric acid as crucial discriminators. Figure 5 also revealed a striking heterogeneity within the metabolic profiles of individuals across the CONTROL, COVID-19, and POST-COVID-19 groups. This heterogeneity is illustrated by the spread and overlap of SHAP value distributions, signifying the varied influence of individual metabolites on the model’s predictions (local explainability). For example, within the CONTROL vs COVID-19 comparison, the spread of data points in the COVID-19 group across higher SHAP values for metabolites like Phenylalanine and Kynurenine indicates a diverse metabolic response to the infection (Figure 5A). Similarly, within the Control vs POST-COVID-19 comparison, the POST-COVID-19 group shows a range of SHAP values for metabolites such as Taurine and LysoPC a C16:0, reflecting the varied trajectories of metabolic recovery or persisting alterations post-infection (Figure 5B). This metabolic diversity underscores the complex non linear relationships and the utility of machine learning models in capturing and interpreting these differences at an individual level.

**Figure 5.**
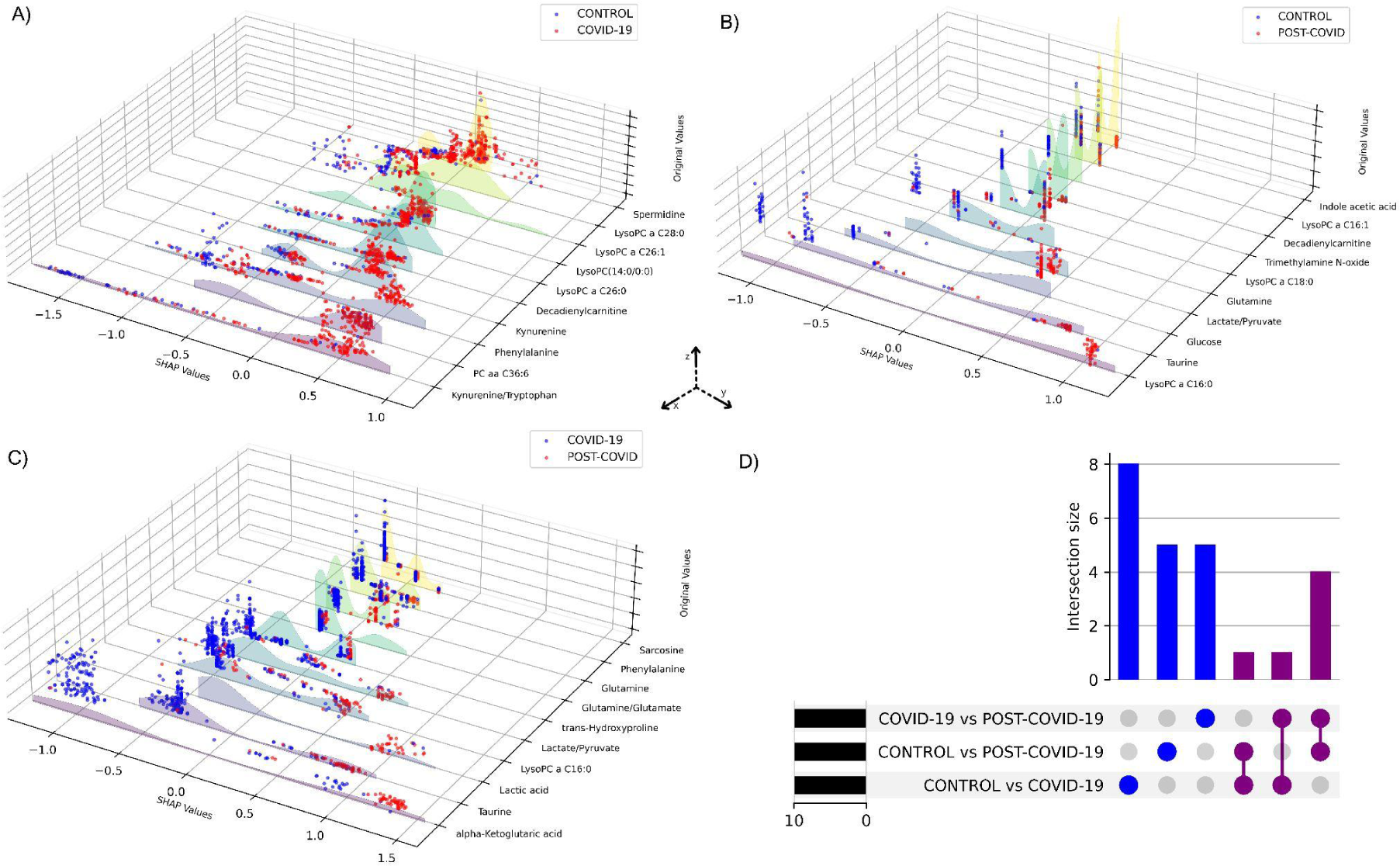
SHAP values of metabolites from binary XGBoost models for distinct group combinations. 3D plots demonstrate SHAP value comparisons between groups: (A) CONTROL vs COVID-19, (B) CONTROL vs POST-COVID-19, and (C) COVID-19 vs POST-COVID-19. Each blue or red dot represents a CONTROL, COVID-19 or POST-COVID-19 according to the respective panel. The y-axis displays the SHAP values, the x-axis ranks the metabolites by their importance to the model and the z-axis shows the metabolites’ original values. Higher SHAP values suggest a greater positive influence on the prediction classification, whereas lower values indicate a negative influence. D) UpSet plot visualizing the intersection of metabolites among three comparisons: CONTROL vs COVID-19, CONTROL vs POST-COVID-19, and COVID-19 vs POST-COVID-19.

### Metabolic subgroup discovery using Explainable embeddings with UMAP and SHAPley values

Building upon the insights gained from the SHAP analysis, which highlighted the specific metabolic influence on our XGBoost model’s predictions and inferable heterogeneity, we explored at a deeper level the metabolic rules that potentially underlie the classes in our data. To achieve a finer-granular metabolic profiles explanation, we utilized a supervised SHAP-based clustering strategy to define a set of decision rules capable of dissecting the local explainability of the data ^30^. Specifically, this task was accomplished through four steps (Figure 1, Section Subgroup discovery). First, we select a pairwise comparison and calculate the SHAP values of all metabolites from a XGBoost model. Then, by considering all the local explanations of all metabolites for all the patients (SHAP values matrix), we visualized their topological structure into a low-dimensional space through UMAP. Posteriorly, in this reduced space, we calculated the number of clusters through HDBSCAN. Finally, having identified the clusters of samples, we trained a multiclass XGBoost model and identified the set of metabolites and their rules to classify each cluster. Then, for the decision rules for each cluster, we used dependence plots to illustrate the concentration of metabolites (original concentration values) versus their corresponding SHAP values, along with clusters. This was done to determine decision rules for operators such as lower than (<), or higher than (>), or its combination (for non-linear interactions) of metabolite concentrations (see Figure 6 and an example in Supplementary Figure 1). We determined the top variables for each cluster until the cluster of interest showed no clear separation in the dependence plot.

**Figure 6.**
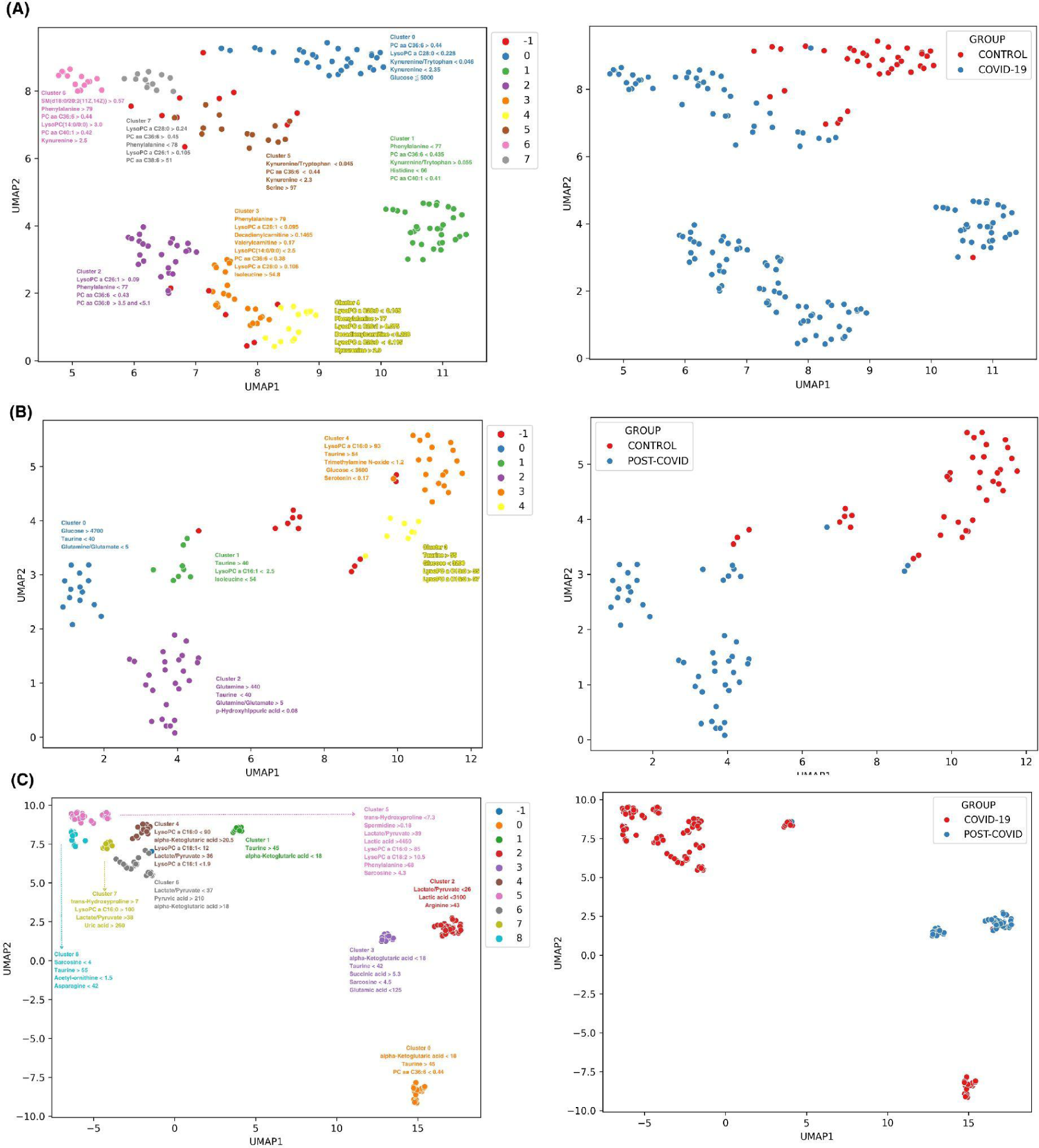
Discrimination and subgroup classification employing UMAP and HDBSCAN on SHAP value distributions from XGBoost models. Each point represents a sample, color-coded by distinct metabolic clusters. A) CONTROL vs COVID-19 with 8 clusters, B) CONTROL VS POST-COVID-19 with 5 clusters, and C) COVID-19 VS POSTCOVID-19 with 9 clusters. Note: Cluster −1 consists of samples that were not dense enough to be classified into a group with HDBSCAN and are classified as noise samples. Clusters are annotated with the most influential features contributing to the subgrouping. The unit of concentration for each numerical value is micromoles as the original data. The right panels shows a color-coded binary classification of samples of their original groups

As a result, several sub-groups of metabolites with similar contributions in the classification were discerned, prompting a deeper understanding of disease progression and its metabolic footprint. This rigorous approach culminated in the derivation of sub-group decision rules (See methods).

Figure 6A shows eight discernible metabolic clusters from CONTROL vs COVID-19. Notably, although certain metabolic markers were predominant in most clusters (such as the kynurenine/tryptophan ratio, kynurenine, and phenylalanine levels, initially identified in our multiclass machine learning model), each cluster features a characteristics combination and concentration of metabolites. CONTROL vs POST-COVID-19 showed a total of five metabolic clusters (Figure 6B). These clusters showcased distinct markers, with some showing pronounced levels of taurine and glucose concentrations, which emphasized the importance of these 2 metabolites. Notably 3 of the 5 clusters were associated with POST-COVID-19, but showed no difference using the number of symptoms(Supplementary Figure 4). For COVID-19 vs POST-COVID-19 (Figure 6C), we observed nine clusters. Metabolites such as alpha-Ketoglutaric acid, taurine, and the lactate/pyruvate ratio characterized the clusters, notably 3 of the 4 clusters characterized from POST-COVID-19 patients showed greater distance from COVID-19 clusters, indicating the heterogeneity of the disease.

Overall, the combination of UMAP and SHAP values for interpretability of the data allowed to draw two main results. First, at higher levels of resolution, there is heterogeneity in the compositions of the samples, even classified in the same clinical group. Second, as expected, each group inside the physiological class has different rules of classification given by concentrations of a few sets of metabolites. These last rules of classification, results substantially contribute to selecting those metabolites for achieving global or local classification with potential application in clinical sets. Thus, our analysis is able to analyze the composition of different physiological groups inside the same patients of COVID-19, and postulate the rules that contribute to their metabolic classifications.

## Discussion

In confronting the monumental health crisis posed by COVID-19 and its sequelae, a thorough understanding of its metabolic implications is paramount. By employing a multi-modal analytical approach, encompassing metabolomics and advanced machine learning techniques, our investigation highlights the complex interplay of metabolites during and post-infection.

The intricate landscape of metabolomic alterations that we observed supports the hypothesis that SARS-CoV-2 infection leads to profound and sustained disruptions in metabolic homeostasis; however, the full alterations panorama cannot be evaluated taking into account just the linear interactions. The use of PCA and PLS-DA initially revealed an overlapping metabolic landscape across CONTROL, COVID-19, and Post-COVID-19 conditions, underscoring the complex nature of metabolic dysregulation that linear methods alone cannot parse. This aligns with emerging evidence of the multi-faceted metabolic response to SARS-CoV-2 that linear models can only partially unravel due to their inability to capture nonlinear topologies ^31^. By UMAP and EMD analyses, we observed clearer demarcations between health states, indicative of the method’s sensitivity to the non-linear interactions of metabolic features. This reinforces the premise that COVID-19’s metabolic disruption cannot be described within a linear framework. It’s particularly noteworthy that while UMAP afforded better group separation, it remained influenced by feature magnitude, a limitation that might obscure metabolites with lower abundance but high biological significance^32^. For instance, the persistence of certain metabolomic imbalances in the Post-COVID-19 phase, underscores the enduring nature of the viral impact, which could potentially inform the etiology of long-lasting symptoms experienced by patients^18^.

XGBoost with SHAP explainability circumvented the pitfalls of magnitude biases and boosted its interpretability. The data analysis discussed here offers a refined metabolic landscape, accentuating subtle yet influential metabolites such as PC aa 36:6 and Taurine across the COVID-19 and Post-COVID-19 states. The emergence of XGBoost’s superior predictive performance, with AUC scores attaining near-perfect metrics, reflects its adeptness at modeling complex, high-dimensional data. This not only validates the algorithm’s application in high-throughput metabolomics data but also demonstrates its potential in clinical settings for prognosticating disease trajectories, such as differentiating states of a healthy state or Post-COVID-19, as its has been proven in other diseases^33–35,36–38^.

By identifying the most influential metabolites in our classifications, SHAP values have highlighted key metabolites that may play crucial roles in the pathogenesis of COVID-19 and Post-COVID-19 syndrome. According to the SHAP values, the disrupted metabolomic profile of acute COVID-19 (see Figure 4B and 5A) is primarily associated with metabolites participating in the immune response and energy metabolism based on our top metabolites found, for example elevated SHAP values for metabolites such as Kynurenine, a by-product of the tryptophan metabolism pathway, suggest an activation of indoleamine 2,3-dioxygenase (IDO) due to inflammation^39^. The role of Kynurenine in COVID-19 has been the subject of various studies. It has been found that serum and saliva levels of Kynurenine and the ratio of Kynurenine to tryptophan (KYN:TRP) may reflect the acute and long-term pathophysiology of SARS-CoV-2, suggesting its potential as a biomarker for diagnosis and monitoring the disease and its therapy ^40,41^, also alterations in carbon and nitrogen metabolism in the liver, indicative of broader changes in energy production pathways during COVID-19 have been observed in moderate and severe cases^42^ Moreover, perturbations in lipid molecules like PC aa C36:6 and LysoPC a C18:2 underscore the potential role of lipid metabolism in the virus infection which has been previously described ^43,44^. These observations are pivotal for understanding the pathophysiological processes at play and may aid in the identification of therapeutic targets that could disrupt the virus’s life cycle or mitigate the inflammatory response.

In the Post-COVID-19, SHAP values indicate a distinct shift in metabolite significance with taurine and glutamine standing out (see Figure 4B and 5B). Persistently altered levels of these amino acids point towards a sustained immune challenge^45^ or a delayed return to homeostatic metabolic function post-infection. The consistent impact of taurine, known for its role in bile salt formation and osmoregulation, may also reflect ongoing oxidative stress and or a lack of cellular detoxification^45,46,47^. Glutamine’s role in supporting immune cell energy requirements could signal a protracted recovery phase where the immune system remains engaged beyond the clearance of the virus ^48,49,50^. Understanding these sustained metabolic changes is critical for developing post-acute care strategies and could be integral in preventing long-term sequelae often observed in Post-COVID-19 syndrome patients. Our findings reinforce the observation that there are metabolic pathways that remain altered even in the post-recovery phase ^17,28^. For instance, persistent fatigue, a hallmark of Long COVID-19^51^, may be tied to the disruptions in energy-related metabolites that we observed, in this instance Taurine supplementation could be used for patients that have lower levels of this metabolite to counter its symptom^52^.

Our Semi-Supervised SHAP-based Clustering strategy (see Figure 6), allowed for the intricate subgroup discovery beyond traditional analytical capacities. This novel methodological approach eschews simple distance metrics, instead emphasizing the discriminative importance of metabolites as determined by their contribution to the model’s predictive accuracy. This machine learning analysis reveals the diversity in the metabolic response to SARS-CoV-2 infection and the varied recovery patterns, which are often homogenized in broader analyses. In the comparison between CONTROL and COVID-19 samples, we observed eight distinct metabolic clusters (see Figure 6A). Each subgroup within the COVID-19 group displayed unique metabolic derangements, indicating the possibility of different viral response phenotypes or stages of disease progression, normally COVID-19 is classified using the WHO classification which ranges from asymptomatic to critical illness^53^, but a more detailed subclassification could be used to improve treatments. The CONTROL vs. Post-COVID-19 (see Figure 6B) analysis presented five metabolic clusters with two key metabolites, taurine and glucose, standing out in their altered levels. The prominence of these metabolites in certain clusters suggests potential pathways that could be investigated for therapeutic interventions. Interestingly, the majority of Post-COVID-19-specific clusters did not correlate with the symptomatology (See Supplementary Figure 4), an observation that points to the complex and possibly non-linear relationship between metabolic alterations and clinical manifestations of Post-COVID-19 syndrome. The variation within Post-COVID-19 clusters indicates possible subtypes of long-term sequelae, underlining the need for personalized approaches in managing these patients. A similar strategy has been applied by Cooper et al^54^ to COVID-19 symptomatology, Cooper identified 16 different clusters of symptoms emphasizing the complex heterogeneity of the disease and the necessity for individualized therapeutic strategies using a holistic approach. For future studies, it will be essential to correlate these metabolic subgroups with clinical outcomes and symptomatology more closely. Prospective studies including longitudinal sampling and in-depth phenotyping are needed to confirm the stability and clinical relevance of these metabolic clusters. Moreover, integrating multi-omics data such as genomics, proteomics, and transcriptomics could offer a systems biology perspective, providing a more comprehensive understanding of the pathophysiological mechanisms at play.

Our study, however, is not without limitations. The reliance on two datasets may introduce biases specific to the population sample. Also, metabolic responses are known to be influenced by a variety of factors including diet, medication, and comorbidities, which were not controlled for, in the datasets. Future research should aim to replicate these findings across diverse cohorts to ensure the generalizability of the metabolic signatures identified for subtyping.

In closing, our investigation offers a robust analytical framework that provides a comprehensive metabolic viewpoint on COVID-19 and its prolonged impact. The application of machine learning models to metabolomics is an approach that holds great promise for elucidating the multifaceted nature of infectious diseases and its long-term consequences. The potential for these findings to inform clinical practice and guide future research is substantial, offering hope for improved management and understanding of Post-COVID-19 syndrome and other similarly complex physiological conditions.

## Methods

### Data Collection

Datasets were sourced from the Mendeley Database at the following URLs:

● Dataset 1 https://data.mendeley.com/datasets/8zfdjsypd8/1
● Dataset 2 https://data.mendeley.com/datasets/7fnt3nfhdv/2.

Lopez et al. employed these datasets in two distinct studies, aiming to pinpoint biomarkers and discern metabolic alterations tied to COVID-19 and its post-infection phase^16,28^. Both investigations utilized an identical method to yield quantitative readings for 108 metabolites, with the omission of carnitine C14:1 in the Post-COVID-19 dataset. To maintain the veracity of the original concentration values, we did not normalize or scale the metabolite data prior to its use in the machine learning models.

### Principal Component Analysis (PCA), Partial Least Square Discriminant Analysis (PLS-DA)

PCA is a statistical technique that transforms the original variables into a new set of uncorrelated variables known as principal components. These components capture the majority of variance present in the original dataset, and in doing so, reveal dominant patterns. PLS-DA, similar in spirit to PCA, is designed to find the direction in the multivariate space that maximizes the separation between classes or groups. It’s particularly suitable for datasets with more variables than observations. PCA and PLS-DA were conducted using the Metaboanalyst 5.0 software^55^. During these analyses, we recognized the need for data normalization to mitigate any artifacts as these techniques perform worse without normalization. As such, we tried multiple normalization procedures to ascertain optimal parameters. The applied normalization techniques encompassed median normalization, log transformation, and pareto scaling. The specific results of these procedures are showcased in the Supplementary Figure 1.

### Differential Expression Analysis via Earth Mover’s Distance (EMD)

EMD offers a way to measure the “distance” between two probability distributions over a region. It can be perceived as the least amount of work needed to transform one distribution into the other. To dissect differential expression in metabolite data, Earth Mover’s Distance (EMD) was used. This method adeptly captures differences in data distributions. The analysis was performed using the “scprep” library in Python, contrasting EMD values across all the datasets. The derived EMD outcomes rendered a ranked inventory of metabolites, underscoring their relative expression shifts. A positive value indicates that transforming the distribution of that metabolite in the group corresponding to the column (COVID-19, CONTROL, POST-COVID) into the distributions of the metabolite in the other groups requires more “work.” Conversely, in general, a positive EMD value means that it is elevated compared to the other groups, while a negative value indicates that it has a decreased value.

### Machine Learning Model Implementation

Different machine learning models were used. The array of machine learning algorithms we tapped into were:

● **Random Forest (RF)**: An ensemble method that constructs multiple decision trees during training and outputs the class that is the mode of the classes for classification, or average prediction for regression.
● **XGBoost**: An optimized gradient boosting library designed to be highly efficient, flexible, and portable.
● **Logistic Regression (LR)**: A regression analysis method suited for prediction of outcome of a categorical dependent variable based on one or more predictor variables.
● **Support Vector Machine (SVM)**: A supervised machine learning algorithm which can be employed for both classification or regression challenges.

Machine learning algorithms were implemented in Google Colab with Python (v. 3.10). Random forest (RF), XGBoost, logistic regression (LR), and support vector machine (SVM) were written using scikit-learn package. To evaluate the performance of the models, the dataset was split into training and testing sets, the training set comprised 80% of the data, while the remaining 20% was allocated for testing.

### Model Evaluation

Each model’s efficacy was estimated using a blend of cross-validation and specific evaluation metrics. A 10-fold cross-validation was executed on the training subset, with accuracy, precision, recall, and F1 score computed through the cross_val_score function from scikit-learn. Post cross-validation, models were further appraised on the testing set using the aforementioned metrics. The superior model was identified based on its performance metrics, encapsulated by:

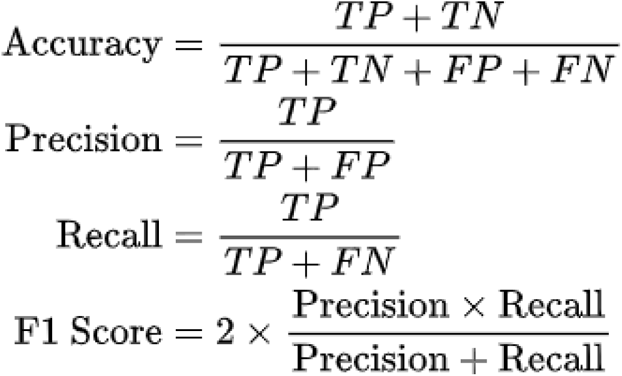

### Hyper-parameter Tuning

In the domain of machine learning, the enhancement of model performance is frequently achieved through a meticulous process termed as hyperparameter optimization. For our research, the optimization strategy employed was using a combination of randomized search and cross-validation methodologies. Randomized search, distinct from the exhaustive nature of grid search, offers an efficient exploration of the hyperparameter space by examining a random subset of possible parameter values, leading to faster convergence to optimal values. Complementing this, cross-validation ensured that the model’s evaluation was robust and unbiased by systematically partitioning the dataset into training and validation subsets. For specificity, the hyperparameters scrutinized for each predictive algorithm were:

1. Random Forest (RF):

○ **n_estimators**: Reflecting the count of trees in the forest, which determines the ensemble’s complexity and predictive capability.
○ **max_depth**: Signifying the maximum number of levels in each decision tree, thereby controlling the depth and potential overfitting.
○ **min_samples_split**: Denoting the minimal count of data points placed in a node before the node is split.
○ **min_samples_leaf**: The minimum number of data points allowed in a leaf node.
2. XGBoost:

○ **n_estimators**: Corresponding to the total count of sequential trees to be modeled.
○ **max_depth**: Dictating how deeply each tree can grow during any boosting round.
○ **learning_rate**: Adjusting the contribution of each tree to the final outcome.
○ **subsample**: The fraction of samples used for fitting the individual base learners.
3. Support Vector Machine (SVM):

○ **C**: Regularization parameter that determines the trade-off between achieving a low margin and ensuring the classifier segments most of the data points correctly.
○ **kernel**: Specifies the type of hyperplane utilized to separate the data.
○ **gamma**: Parameter for non-linear hyperplanes, determining the curve’s fit to the data.
4. Logistic Regression:

○ **C**: Inverse regularization strength, which can prevent potential overfitting.
○ **penalty**: Denoting the norm utilized in the penalization.
○ **solver**: Algorithmic approach employed for optimization problems.

### Shapley Values

Shapley Additive exPlanations (SHAP) facilitates local prediction interpretations by ascertaining the importance of each metabolomic feature per sample prediction. As a robust post-hoc IML method, SHAP extends comprehensive global model insights. Rooted in the cooperative game theory methodology of Shapley values^26,54^, SHAP offers a fair approach to apportion rewards within a cooperative game. In this context, the game represents the machine learning model, and the Shapley value fairly ascribes each metabolomic feature’s contribution to the outcome. We computed the Shapley values via the shapTreeExplainer, using the Python SHAP package.

### Supervised clustering using Local Explanations and Manifold Learning

Binary XGBoost models were trained for binary classification, integrating SHAP values for supervised clustering. Post-hoc analysis with SHAP and nonlinear dimensionality reduction using UMAP provided subgroup characterizations within SHAP values data. Hierarchical Density-Based Spatial Clustering of Applications with Noise (HDBSCAN) identified samples with similar metabolite classification weights. Decision rules were elucidated via a dependency plot, mapping SHAP values against significant metabolites for each cluster. To establish the decision rules for each cluster, we utilized dependence plots that depict the concentration of metabolites (original values) against their corresponding SHAP values, accompanied by clusters. This approach enabled us to formulate decision rules involving operators such as <, >, or combinations thereof (for non-linear interactions) concerning metabolite concentrations. We continued to identify the most significant variables for each cluster until no clear separation was observable in the dependence plot for the cluster of interest.

### Code

All codes used in this study are available at https://github.com/resendislab/POST_COVID_Metabolome_MachineLearning

## Acknowledgements

JJOV work was supported by CONACYT (Grant Ciencia de Frontera 2019, FORDECYT-PRONACES/425859/2020) and UNAM Posdoctoral Program DGAPA (POSDOC). C-PM. is a doctoral student from Programa de Doctorado en Ciencias Biomédicas, Universidad Nacional Autónoma de México (UNAM) and received fellowship to CVU 855825 from CONAHCyT, México.

## Supplementary information

**Supplementary Figure 1:**
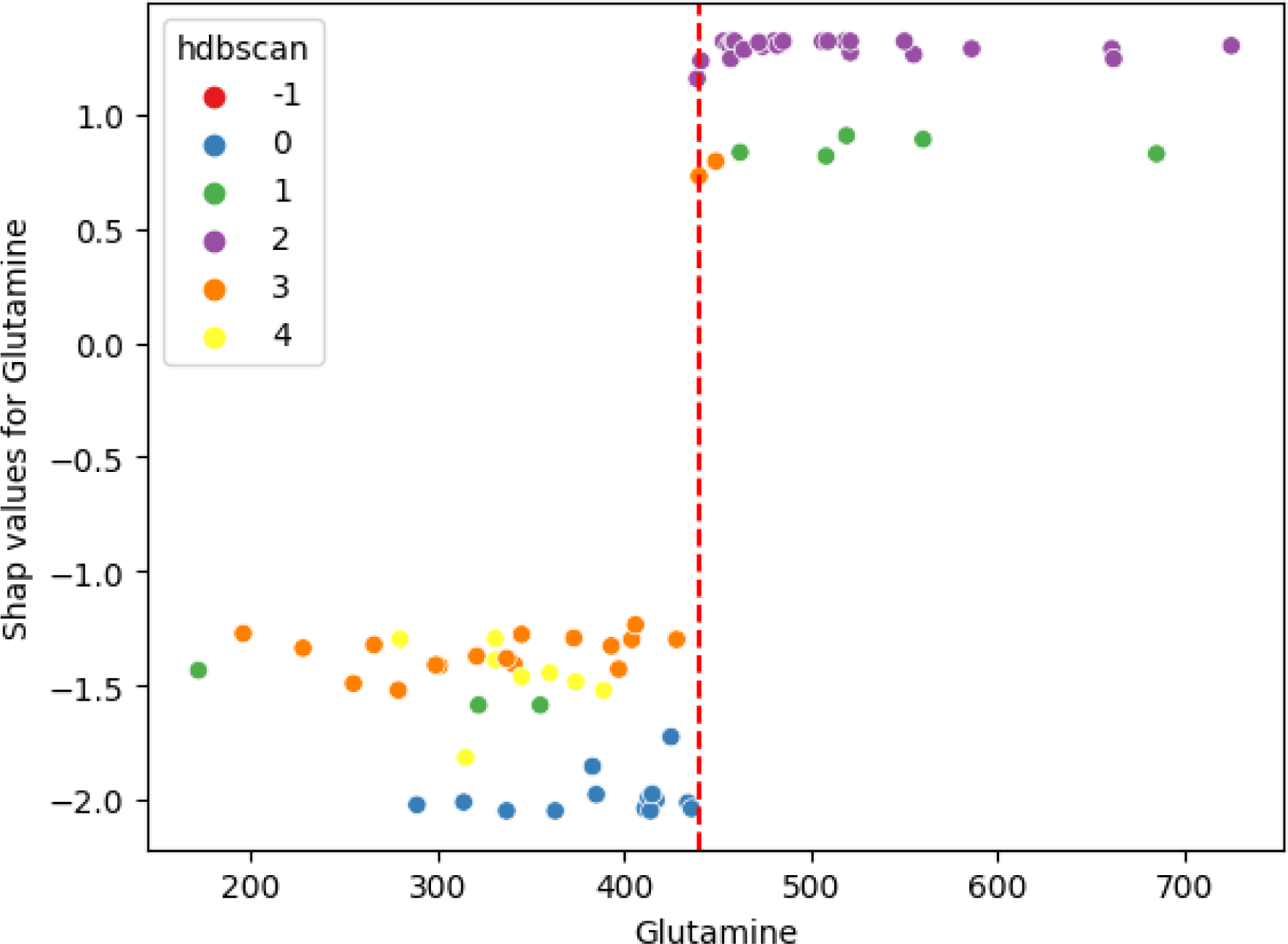
Dependency Plot for Cluster 2 in the Control vs. Post-COVID Comparison. The scatter plot displays the relationship between glutamine levels and their corresponding SHAP values for Cluster 2, as identified by HDBSCAN. Each point represents an individual sample, color-coded according to its HDBSCAN subgroup assignment. The vertical dashed red line indicates the threshold level of glutamine used as its decision rules.

**Supplementary Figure 2.**
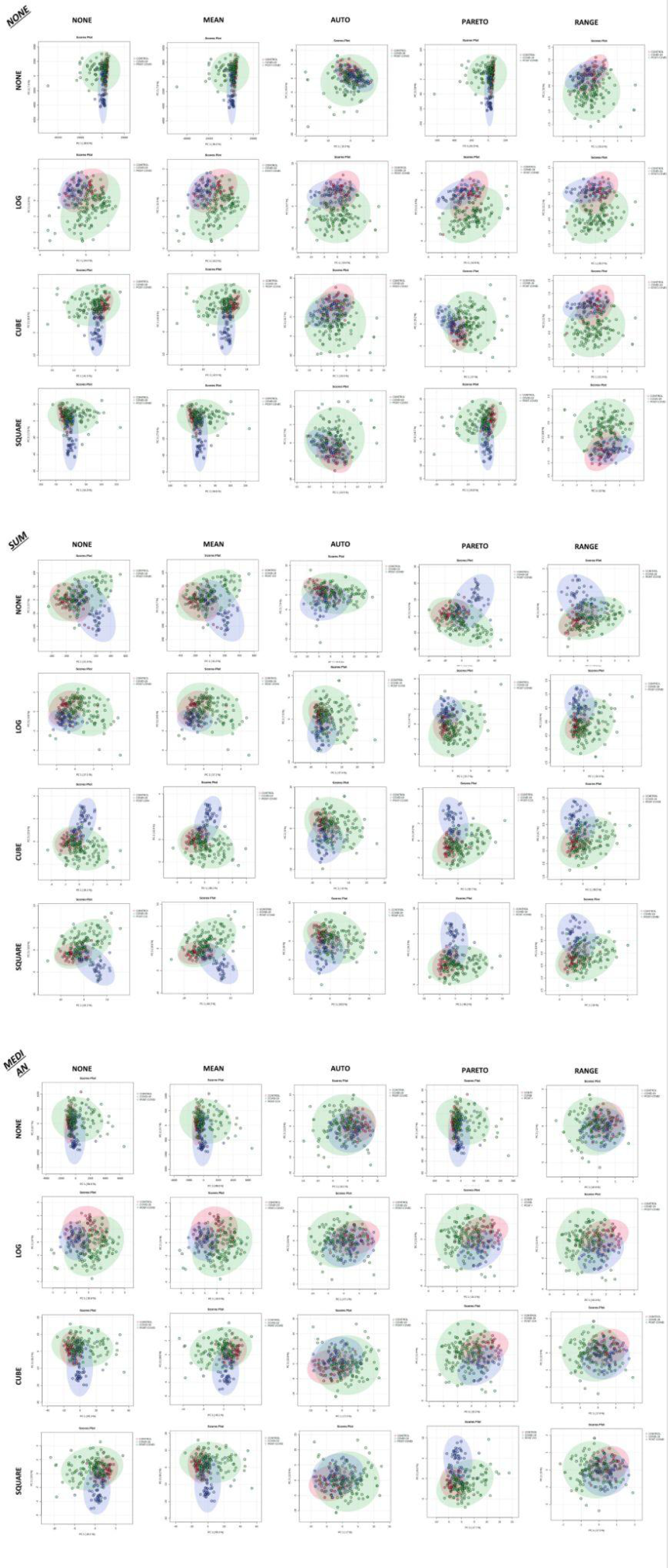
Multiple PCA plots using different normalization strategies. Each panel shows a PCA score plot of the same dataset, but normalized using a different strategy. Top left corner shows the normalization strategy (None, Sum, Median), the top row shows the different types of scaling (None, Mean, Auto, Pareto and Range), the left column shows transformation strategies with square root, cube root, and logarithmic transformations.

**Supplementary table 1.** Performance metrics of machine learning binary models.

|  |  |  | Cross Validations (CV) |  |  |  |  |  |
| --- | --- | --- | --- | --- | --- | --- | --- | --- |
| Comparisons | AUC | Accuracy | CV 1 | CV 2 | CV 3 | CV 4 | CV 5 | Average CV |
| **CONTROL VS COVID-19** | 0.991 | 0.944 | 0.861 | 0.889 | 0.889 | 0.972 | 0.917 | 0.906 |
| **CONTROL vs POST-COVID** | 1.000 | 0.944 | 1.000 | 1.000 | 1.000 | 1.000 | 0.882 | 0.976 |
| **COVID-19 VS POST-COVID** | 0.983 | 0.921 | 0.947 | 0.921 | 1.000 | 0.974 | 1.000 | 0.968 |

**Supplementary table 2.** Top 10 features based on SHAP values in the binary machine learning model XGBoost.

| **CONTROL vs COVID-19** | **CONTROL vs POST-COVID-19** | **COVID-19 vs POST-COVID-19** |
| --- | --- | --- |
| Kynurenine/Tryptophan | LysoPC a C16:0 | alpha-Ketoglutaric acid |
| PC aa C36:6 | Taurine | Taurine |
| Phenylalanine | Glucose | Lactic acid |
| Kynurenine | Lactate/Pyruvate | LysoPC a C16:0 |
| Decadienylcarnitine | Glutamine | Lactate/Pyruvate |
| LysoPC a C26:0 | LysoPC a C18:0 | trans-Hydroxyproline |
| LysoPC(14:0/0:0) | Trimethylamine N-oxide | Glutamine/Glutamate |
| LysoPC a C26:1 | Decadienylcarnitine | Glutamine |
| LysoPC a C28:0 | LysoPC a C16:1 | Phenylalanine |
| Spermidine | Indole acetic acid | Sarcosine |

**Supplementary Figure 3:**
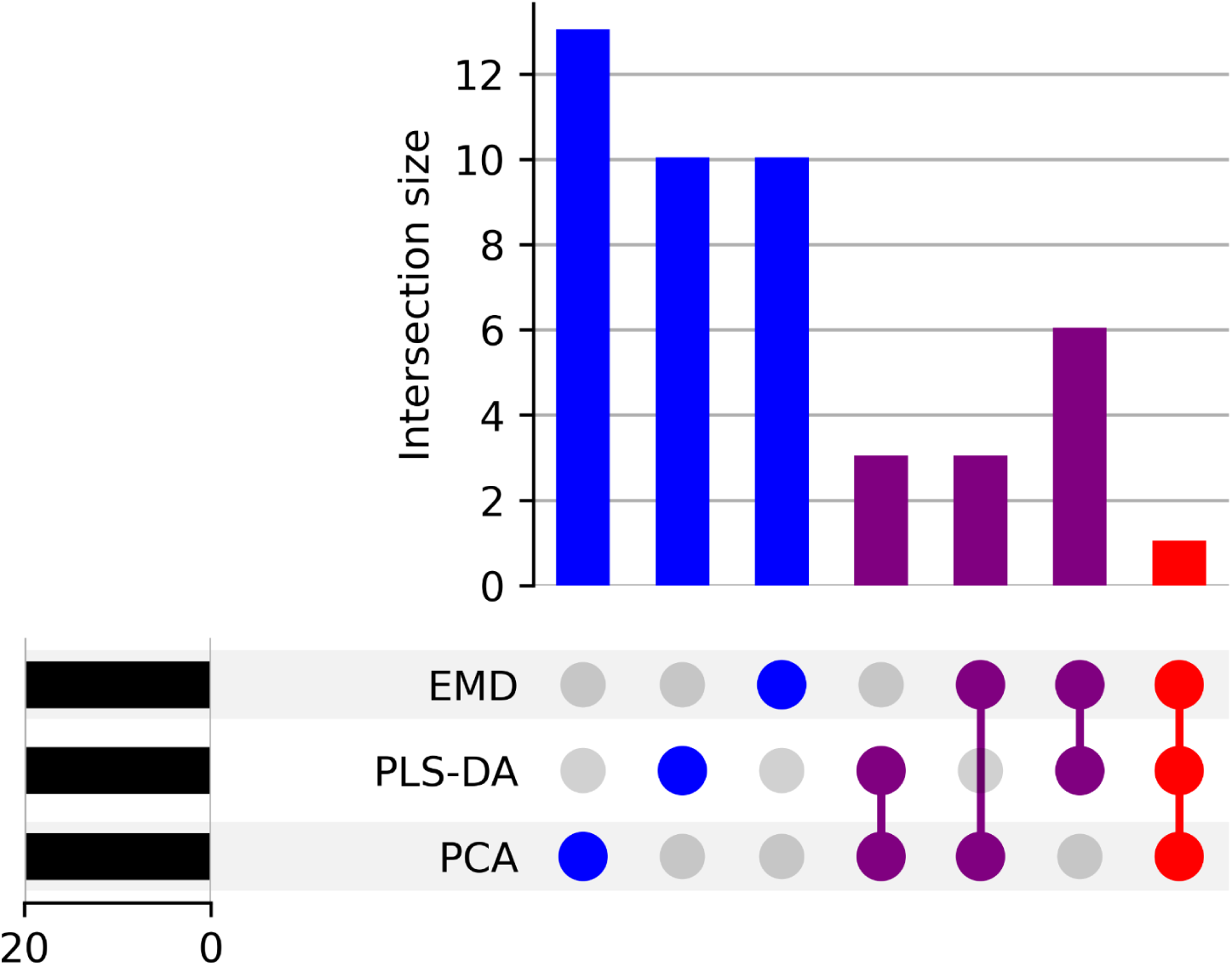
UpSet plot illustrating the intersection sizes and relationships of the top 20 metabolites identified by three different statistical methods. Bar chart represents the number of shared metabolites between the methods. The matrix below shows the presence (colored dot) or absence (empty circle) of a particular method’s metabolites in each intersection. The methods compared are Principal Component Analysis (PCA), Partial Least Squares-Discriminant Analysis (PLS-DA), and Earth Mover’s Distance (EMD). Metabolites for PCA were selected based on loadings from PC1, for PLS-DA by a Variable Importance in Projection (VIP) score greater than 1.5, and for EMD by their absolute z-score differences. The horizontal bars indicate the number of metabolites identified by each method individually, while the vertical bars represent the size of the intersection sets.

**Supplementary Figure 4.**
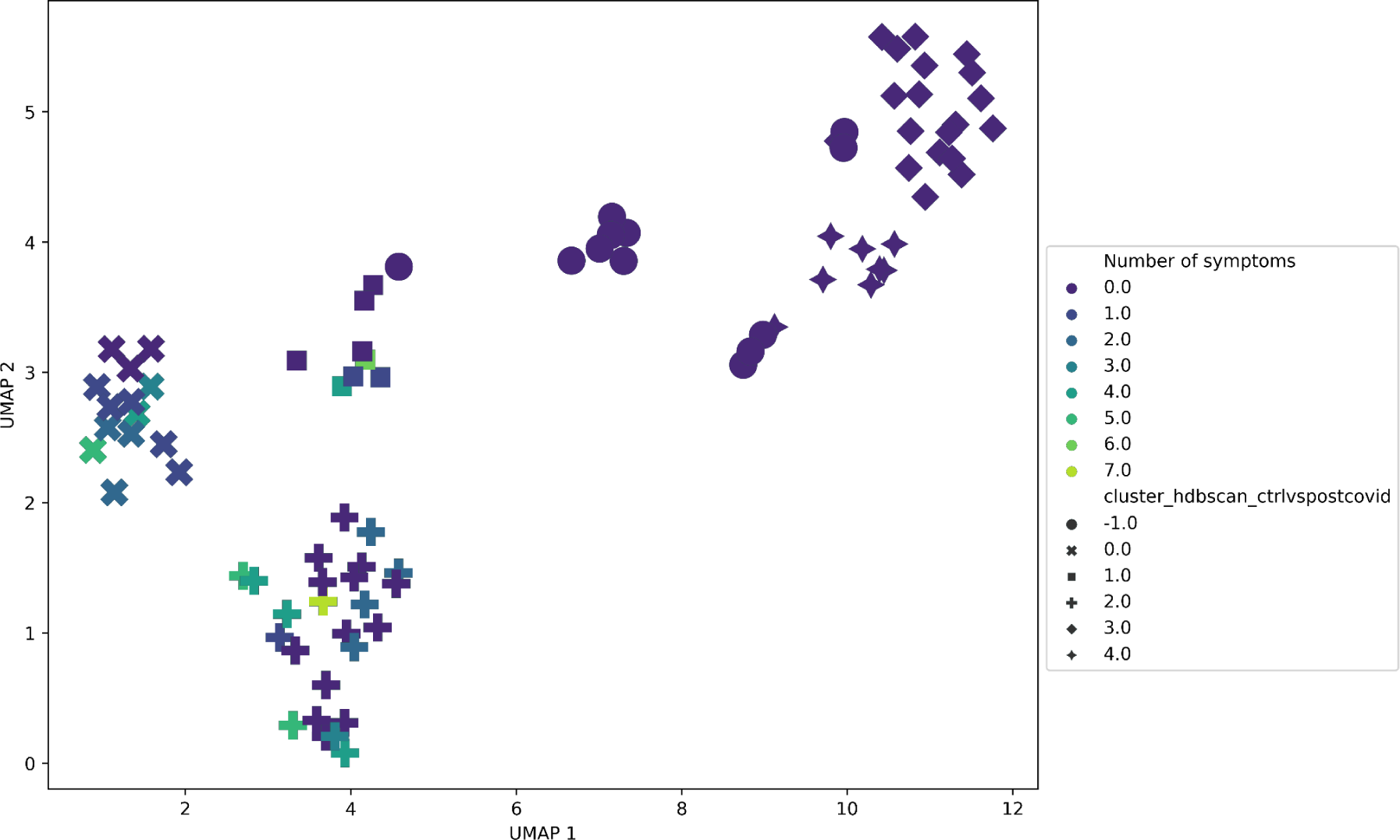
Scatter plot of UMAP-based visualization of patients with POST-COVID, colored by the number of reported symptoms and symbol by HDBSCAN cluster number. The UMAP coordinates were generated using SHAP values from a binary machine learning model CONTROL vs POST-COVID.

## References

1. WHO Coronavirus (COVID-19) Dashboard. https://covid19.who.int/?mapFilter=deaths.

2. Zhao, W., Li, H., Li, J., Xu, B. & Xu, J. The mechanism of multiple organ dysfunction syndrome in patients with COVID-19. J. Med. Virol. 94, 1886 (2022).

3. Al Sulaiman, K. et al. The clinical outcomes of COVID-19 critically ill patients co-infected with other respiratory viruses: a multicenter, cohort study. BMC Infect. Dis. 23, (2023).

4. Reyes, L. F. et al. Clinical characteristics, risk factors and outcomes in patients with severe COVID-19 registered in the International Severe Acute Respiratory and Emerging Infection Consortium WHO clinical characterisation protocol: a prospective, multinational, multicentre, observational study. ERJ Open Research 8, (2022).

5. Khodeir, M. M. et al. COVID-19: Post-recovery long-term symptoms among patients in Saudi Arabia. PLoS One 16, (2021).

6. Galván-Tejada, C. E. et al. Persistence of COVID-19 Symptoms after Recovery in Mexican Population. Int. J. Environ. Res. Public Health 17, (2020).

7. Phetsouphanh, C. et al. Immunological dysfunction persists for 8 months following initial mild-to-moderate SARS-CoV-2 infection. Nat. Immunol. 23, 210–216 (2022).

8. CDC. Long COVID or Post-COVID Conditions. Centers for Disease Control and Prevention https://www.cdc.gov/coronavirus/2019-ncov/long-term-effects/index.html (2023).

9. Ballouz, T. et al. Recovery and symptom trajectories up to two years after SARS-CoV-2 infection: population based, longitudinal cohort study. BMJ 381, (2023).

10. Chen, B., Julg, B., Mohandas, S., Bradfute, S. B. & RECOVER Mechanistic Pathways Task Force. Viral persistence, reactivation, and mechanisms of long COVID. Elife 12, (2023).

11. Iqbal, P., et al. Post-COVID-19-associated multiorgan complications or ‘long COVID’ with literature review and management strategy discussion: A meta-analysis. Health Science Reports 6, (2023).

12. Chen, P., Wu, M., He, Y., Jiang, B. & He, M.-L. Metabolic alterations upon SARS-CoV-2 infection and potential therapeutic targets against coronavirus infection. Signal Transduction and Targeted Therapy 8, 1–23 (2023).

13. Metabolomics: Strategies to Define the Role of Metabolism in Virus Infection and Pathogenesis. in Advances in Virus Research vol. 98 57–81 (Academic Press, 2017).

14. Palmer, C. S. Innate metabolic responses against viral infections. Nature Metabolism 4, 1245–1259 (2022).

15. Rahman, M. & Schellhorn, H. E. Metabolomics of infectious diseases in the era of personalized medicine. Front. Mol. Biosci. 10, 1120376 (2023).

16. López-Hernández, Y. et al. Targeted metabolomics identifies high performing diagnostic and prognostic biomarkers for COVID-19. Sci. Rep. 11, 14732 (2021).

17. López-Hernández, Y. et al. Untargeted analysis in post-COVID-19 patients reveals dysregulated lipid pathways two years after recovery. Front Mol Biosci 10, 1100486 (2023).

18. Liptak, P. et al. Persistence of Metabolomic Changes in Patients during Post-COVID Phase: A Prospective, Observational Study. Metabolites 12, (2022).

19. Tebani, A., Afonso, C. & Bekri, S. Advances in metabolome information retrieval: turning chemistry into biology. Part II: biological information recovery. J. Inherit. Metab. Dis. 41, 393–406 (2018).

20. Ruiz-Perez, D., Guan, H., Madhivanan, P., Mathee, K. & Narasimhan, G. So you think you can PLS-DA? BMC Bioinformatics 21, 1–10 (2020).

21. Shiokawa, Y., Date, Y. & Kikuchi, J. Application of kernel principal component analysis and computational machine learning to exploration of metabolites strongly associated with diet. Sci. Rep. 8, 3426 (2018).

22. McInnes, L., Healy, J. & Melville, J. UMAP: Uniform Manifold Approximation and Projection for Dimension Reduction. (2018) doi:10.48550/ARXIV.1802.03426.

23. Chen, T. & Guestrin, C. XGBoost. in Proceedings of the 22nd ACM SIGKDD International Conference on Knowledge Discovery and Data Mining (ACM, New York, NY, USA, 2016). doi:10.1145/2939672.2939785.

24. Stamate, D. et al. A metabolite-based machine learning approach to diagnose Alzheimer-type dementia in blood: Results from the European Medical Information Framework for Alzheimer disease biomarker discovery cohort. Alzheimers. Dement. 5, 933–938 (2019).

25. Lundberg, S. M. et al. From Local Explanations to Global Understanding with Explainable AI for Trees. Nat Mach Intell 2, 56–67 (2020).

26. Lundberg, S. & Lee, S.-I. A unified approach to interpreting model predictions. (2017) doi:10.48550/ARXIV.1705.07874.

27. Bifarin, O. O. Interpretable machine learning with tree-based shapley additive explanations: Application to metabolomics datasets for binary classification. PLoS One 18, e0284315 (2023).

28. López-Hernández, Y. et al. The plasma metabolome of long COVID patients two years after infection. Sci. Rep. 13, 12420 (2023).

29. Evans, E. D. et al. Predicting human health from biofluid-based metabolomics using machine learning. Sci. Rep. 10, 17635 (2020).

30. Chmiel, F. P. et al. Using explainable machine learning to identify patients at risk of reattendance at discharge from emergency departments. Sci. Rep. 11, 21513 (2021).

31. van der Maaten, L., Postma, E. O. & van den Herik, J. Dimensionality Reduction: A Comparative Review. Tilburg University Technical Report **TiCC-TR**, 2009–2005 (2009).

32. Ghojogh, B., Ghodsi, A., Karray, F. & Crowley, M. Uniform Manifold Approximation and Projection (UMAP) and its variants: Tutorial and survey. (2021) doi:10.48550/ARXIV.2109.02508.

33. Guan, X. et al. Construction of the XGBoost model for early lung cancer prediction based on metabolic indices. BMC Med. Inform. Decis. Mak. 23, 1–16 (2023).

34. Roberts, I. et al. Untargeted metabolomics of COVID-19 patient serum reveals potential prognostic markers of both severity and outcome. Metabolomics 18, 6 (2021).

35. Hogan, C. A. et al. Nasopharyngeal metabolomics and machine learning approach for the diagnosis of influenza. EBioMedicine 71, 103546 (2021).

36. Yi, F. et al. XGBoost-SHAP-based interpretable diagnostic framework for alzheimer’s disease. BMC Med. Inform. Decis. Mak. 23, 137 (2023).

37. Moore, A. & Bell, M. XGBoost, A Novel Explainable AI Technique, in the Prediction of Myocardial Infarction: A UK Biobank Cohort Study. Clin. Med. Insights Cardiol. (2022) doi:10.1177/11795468221133611.

38. Cao, Y., Forssten, M. P., Sarani, B., Montgomery, S. & Mohseni, S. Development and Validation of an XGBoost-Algorithm-Powered Survival Model for Predicting In-Hospital Mortality Based on 545,388 Isolated Severe Traumatic Brain Injury Patients from the TQIP Database. J Pers Med 13, (2023).

39. Kim, S., Miller, B. J., Stefanek, M. E. & Miller, A. H. Inflammation-induced activation of the indoleamine 2,3-dioxygenase pathway: Relevance to cancer-related fatigue. Cancer 121, 2129–2136 (2015).

40. Bizjak, D. A. et al. Kynurenine serves as useful biomarker in acute, Long- and Post-COVID-19 diagnostics. Front. Immunol. 13, 1004545 (2022).

41. Cihan, M. et al. Kynurenine pathway in Coronavirus disease (COVID-19): Potential role in prognosis. J. Clin. Lab. Anal. 36, e24257 (2022).

42. Caterino, M. et al. The Serum Metabolome of Moderate and Severe COVID-19 Patients Reflects Possible Liver Alterations Involving Carbon and Nitrogen Metabolism. Int. J. Mol. Sci. 22, (2021).

43. Dissecting lipid metabolism alterations in SARS-CoV-2. Prog. Lipid Res. 82, 101092 (2021).

44. Seitz, T., Setz, C., Rauch, P., Schubert, U. & Hellerbrand, C. Lipid Accumulation in Host Cells Promotes SARS-CoV-2 Replication. Viruses 15, 1026 (2023).

45. Cruzat, V., Macedo Rogero, M., Noel Keane, K., Curi, R. & Newsholme, P. Glutamine: Metabolism and Immune Function, Supplementation and Clinical Translation. Nutrients 10, 1564 (2018).

46. Baliou, S. et al. Protective role of taurine against oxidative stress (Review). Mol. Med. Rep. 24, 1–19 (2021).

47. Thirupathi, A., Pinho, R. A., Baker, J. S., István, B. & Gu, Y. Taurine Reverses Oxidative Damages and Restores the Muscle Function in Overuse of Exercised Muscle. Front. Physiol. 11, 582449 (2020).

48. Glutamine addiction in virus-infected mammalian cells: A target of the innate immune system? Med. Hypotheses 153, 110620 (2021).

49. Aydın, H. et al. Glutamine-Driven Metabolic Adaptation to COVID-19 Infection. Indian J. Clin. Biochem. 38, 83–93 (2022).

50. Shah, A. M., Wang, Z. & Ma, J. Glutamine Metabolism and Its Role in Immunity, a Comprehensive Review. Animals 10, 326 (2020).

51. What Do I Need to Know About Long-Covid-related Fatigue, Brain Fog, and Mental Health Changes? Arch. Phys. Med. Rehabil. 104, 996–1002 (2023).

52. Kim, S.-H. et al. A Comparative Study of Antifatigue Effects of Taurine and Vitamin C on Chronic Fatigue Syndrome. Pharmacology & Pharmacy 13, 300–312 (2022).

53. Clinical Spectrum. COVID-19 Treatment Guidelines https://www.covid19treatmentguidelines.nih.gov/overview/clinical-spectrum/.

54. Cooper, A., Doyle, O. & Bourke, A. Supervised clustering for subgroup discovery: An application to COVID-19 symptomatology. in Communications in Computer and Information Science 408–422 (Springer International Publishing, Cham, 2021).

55. Pang, Z. et al. MetaboAnalyst 5.0: narrowing the gap between raw spectra and functional insights. Nucleic Acids Res. 49, W388–W396 (2021).

