## Supplementary figures and images for "Exploring Metabolic Anomalies in COVID-19 and Post-COVID-19: A Machine Learning Approach with Explainable Artificial Intelligence"

### Supplementary Figure 1

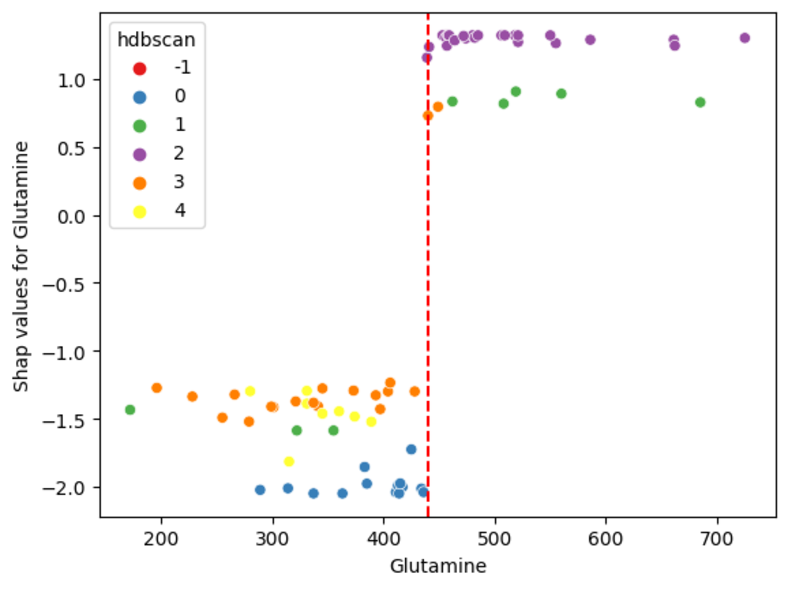

### Supplementary Figure 2

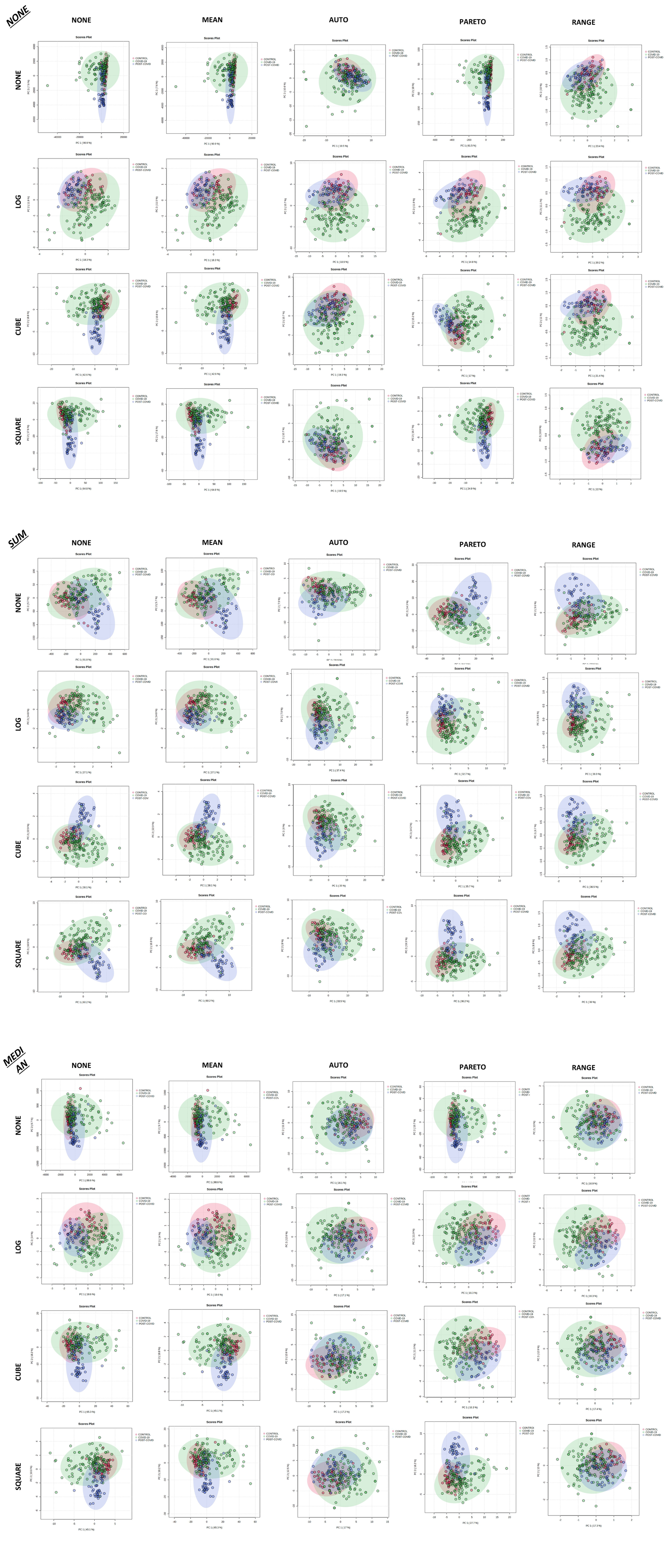

### Supplementary Figure 3

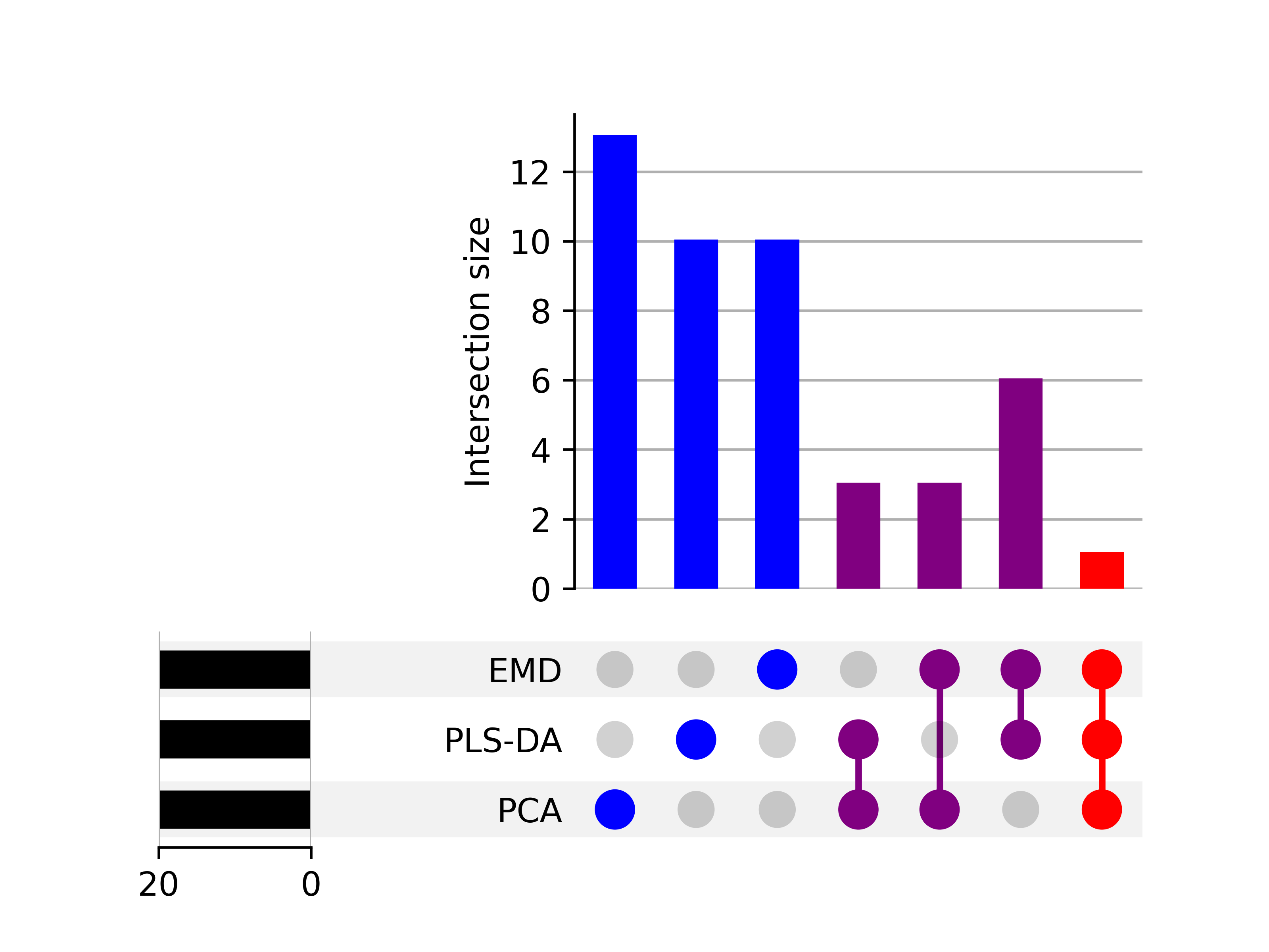

### Supplementary Figure 4

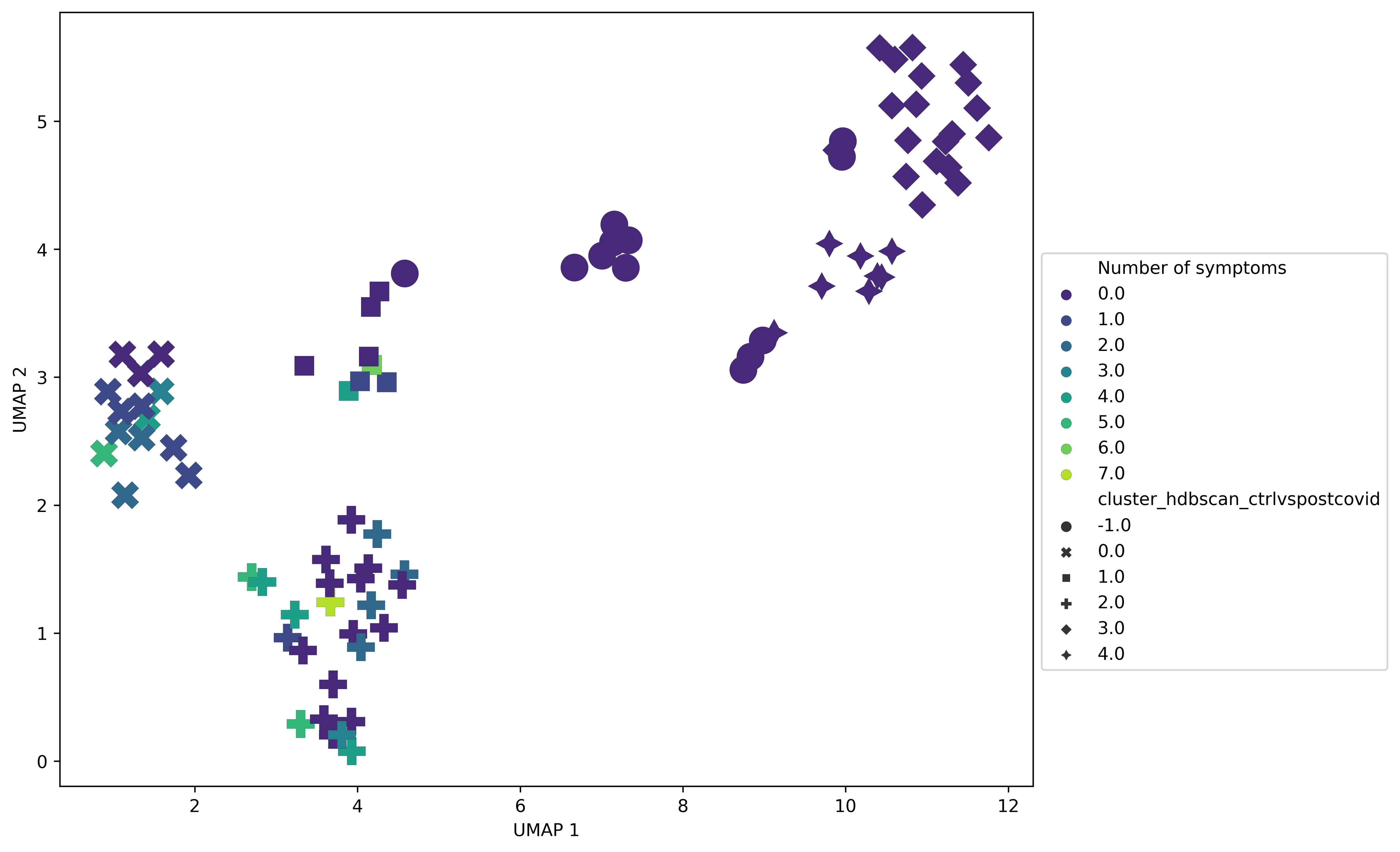
